# Grazer behavior can regulate large-scale patterning of community states

**DOI:** 10.1101/722215

**Authors:** Vadim A. Karatayev, Marissa L. Baskett, David J. Kushner, Nicholas T. Shears, Jennifer E. Caselle, Carl Boettiger

## Abstract

Ecosystem patterning can arise from environmental heterogeneity, biological feedbacks that produce multiple persistent ecological states, or their interaction. One source of feed-backs is density-dependent changes in behavior that regulates species interactions. By fitting state-space models to large-scale (∼500km) surveys on temperate rocky reefs, we find that behavioral feedbacks best explain why kelp and urchin barrens form either reef-wide patches or local mosaics. Best-supported models in California include feedbacks where starvation intensifies grazing across entire reefs create reef-scale, alternatively stable kelp- and urchin-dominated states (32% of reefs). Best-fitting models in New Zealand include the feedback of urchins avoiding dense kelp stands that can increase abrasion and predation risk, which drives a transition from shallower urchin-dominated to deeper kelp-dominated zones, with patchiness at 3-8m depths with intermediate wave stress. Connecting locally-studied processes with region-wide data, we highlight how behavior can explain community patterning and why some systems exhibit community-wide alternative stable states.

## Introduction

Spatial patterning in community types characterizes many ecosystems. For example, in arid ecosystems patches of shrubs and barren soil 25 m in diameter form mosaic patterns (Klausmeier 1999). On mountain ranges, strips of ribbon forests 500m wide intersperse with wider bands of grassy meadows (Hiemstra *et al*. 2006). A longstanding focus of empirical and theoretical work has aimed to resolve the processes generating ecosystem patterns and their spatial scale.

Spatial patterning may occur due to underlying environmental heterogeneity, local biological feedbacks, or an interactive combination of both drivers (Rietkerk & Van de Koppel 2008). Examples of environment-driven patterns include gradients in desiccation stress that zone intertidal communities from heat-tolerant to more sensitive species with increasing water depth (Dayton 1971). Patterning can also arise when biological feedbacks reinforce distinct ecological states. If interactions change from positive to negative with distance between individuals, self-organized patterns of populated and empty areas occur in homogeneous environments, as seen in shrub mosaics in arid ecosystems (Rietkerk & Van de Koppel 2008). Feedbacks that do not change sign with distance can also drive patterns over large scales by amplifying the effects of ecosystem heterogeneity. For instance, grasses predominate in areas of low-moderate herbivory by diluting grazing across many plants while areas with elevated livestock densities exhibit a disproportional plant density collapse via overgrazing (Noy-Meir 1975). In all cases feedback-induced patterning requires strong and often nonlinear species interactions.

One possible driver of biological feedbacks is density-dependent changes in behavior (Peckarsky *et al*. 2008). Herbivory can decline in the presence of predators (McPeek & Peckarsky 1998) or increase when herbivores form large groups that reduce predation risk (Gil *et al*. 2018). On coral reefs, for instance, activity of herbivorous fishes as part of schools can account for 68% of total consumption (Gil & Hein 2017) and influence the potential for coral-dominated or algal-dominated community states (Gil *et al*. 2020). Density-dependent changes in behavior could also create biological feedbacks, for example when dense plant aggregations increase predation risk and decrease herbivory rates, feeding back to increase local plant recruitment. This forms one possible mechanism for why plant stands expand after predator re-introduction (e.g., wolves in Yellowstone, Fortin *et al*. 2005) and recede around refugia from predation following herbivore recovery in the intertidal (Matassa & Trussell 2011) and on coral reefs (Madin *et al*. 2019). However, because isolating the effects of behavior from herbivore density is challenging over large scales (e.g., direct *versus* indirect predator effects, Creel & Christianson 2009), comparing models with and without behavior can provide the next step towards understanding whether behavior can underpin community patterning.

Temperate rocky reefs exemplify each of patterned communities, behavior-mediated herbivory, and environmental variation. Patterning of two distinct ecological states in these ecosystems, kelp forests and urchin-dominated barrens (Fig. 1a, c), can occur at drastically different scales in different regions. Whereas large-scale (> 1km) barrens and forests span all depths on California reefs (Fig. 1d; Cavanaugh *et al*. 2014), in New Zealand meter-scale patchiness occurs at intermediate depths while kelp occupy deeper water and urchins shallower zones (Fig. 1b; Parsons *et al*. 2004). This difference in patchiness scales might arise from a greater sensitivity of the dominant kelp species in New Zealand to wave stress in shallow areas (Grace 1983) and more intensive, larger-scale pulses of urchin recruitment (Hart & Scheibling 1988) or mortality (Lafferty 2004) in California.

**Figure 1:**
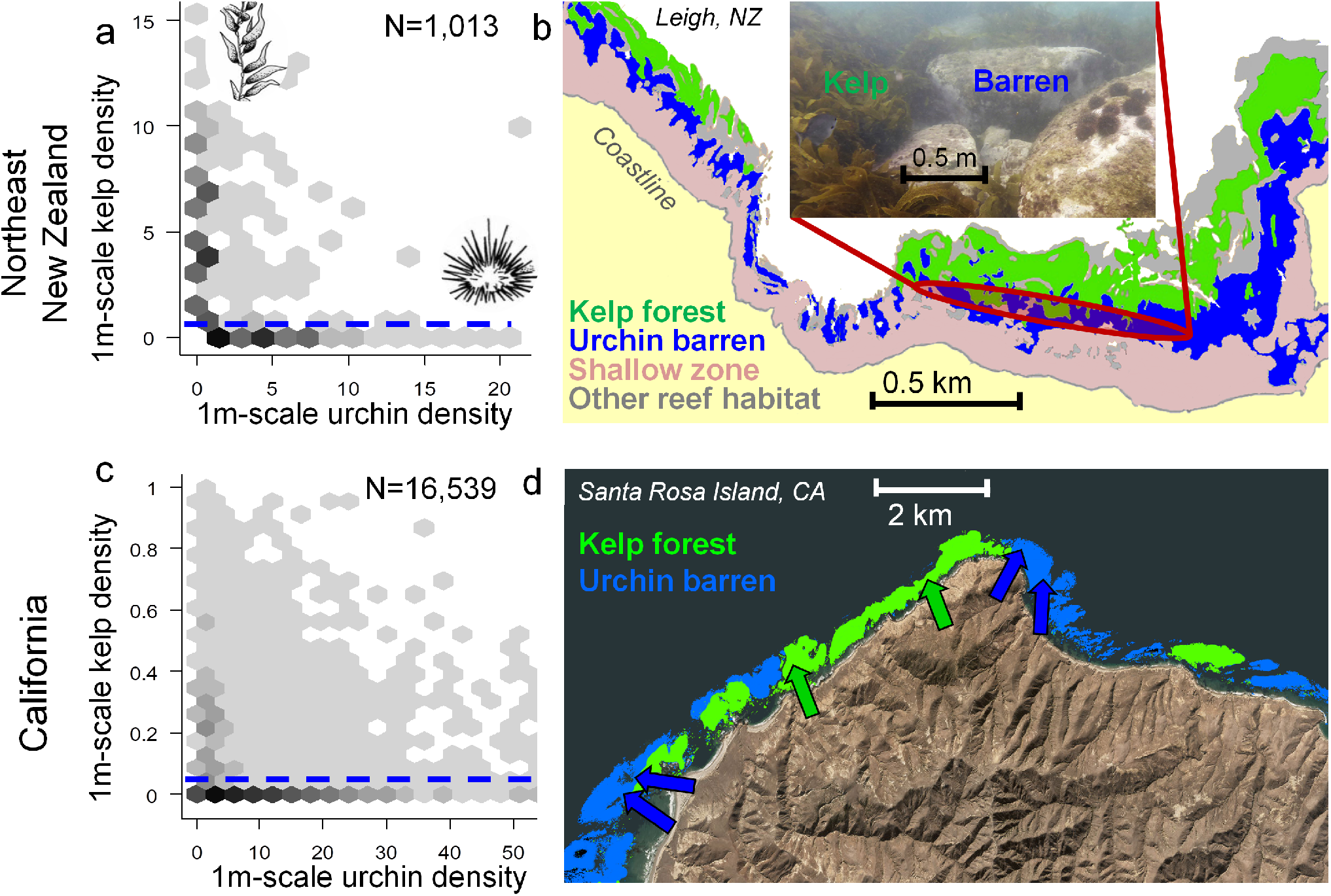
Visualization of kelp and urchin density in New Zealand (a, b) and California (c, d). (a, c) Region-specific distribution of kelp- and urchin-dominated states with N indicating total numbers of samples. Dashed lines in (a, c) denote kelp threshold densities used to categorize community regimes as forested (i.e., exceeding the 10th quantile of kelp densities in samples with kelp in each region) and barren otherwise. (b) Depth zonation of New Zealand reef communities two years after establishment of the Leigh Marine reserve, reproduced from Leleu *et al*. 2012. Inset in shows the local kelp and barren patchiness in mid-depth zones, and “shallow zone” denotes intertidal and shallow sub-tidal areas dominated by wave-tolerant brown algal species other than *Ecklonia*. (d) Reef-scale patchiness of forest and barren community states at Santa Rosa Island, California in 2016 from satellite imagery (shaded areas) and 6 monitoring sites (arrows, with arrow color denoting site state). See Appendix A for details on state estimation from satellite imagery and density measurements in (a, c). Note that exact state classifications may differ between panels (a) *vs*. (b) and (c) *vs*. (d).

Regional differences in temperate reef patchiness could alternatively (or additionally) arise from differences in urchin behavior. Urchins can exhibit two feeding modes: passive grazing on kelp fronds detached from plants and carried to the bottom by currents (‘drift kelp’ hereafter), typically while occupying cryptic habitats (e.g., crevices) protected from predators and physical stress, and active grazing on open substrate (Dayton 1985b; Harrold & Reed 1985). In New Zealand, urchins switch to active grazing in local 1-5m^2^ patches when subcanopy kelp densities become insufficient to deter active grazing via physical abrasion (kelp ‘whiplash’ effects, Konar 2000) or by attracting urchin predators (Cowen 1983; Fig. 2e). In California, giant kelp concentrate biomass in large canopies well above the seafloor, reducing local abrasion while increasing reef-wide drift kelp subsidies. A switch to active grazing can therefore happen synchronously across each reef in California when low kelp densities and low drift kelp subsidies to the seafloor cause reef-wide urchin starvation (Ebeling *et al*. 1985; Harrold & Reed 1985; Fig. 2d). As declines in kelp density increase grazing activity locally (in New Zealand) or reef-wide (in California) and feed back to further deplete kelp, we hypothesize that behavior can create region-specific community patterning. The ability of behavioral feedbacks and distinct states to explain temperate reef spatial patterning also informs whether urchin barrens and kelp forests occur as alternative stable states, a long-debated phenomenon in rocky temperate reefs (Petraitis & Dudgeon 2004).

**Figure 2:**
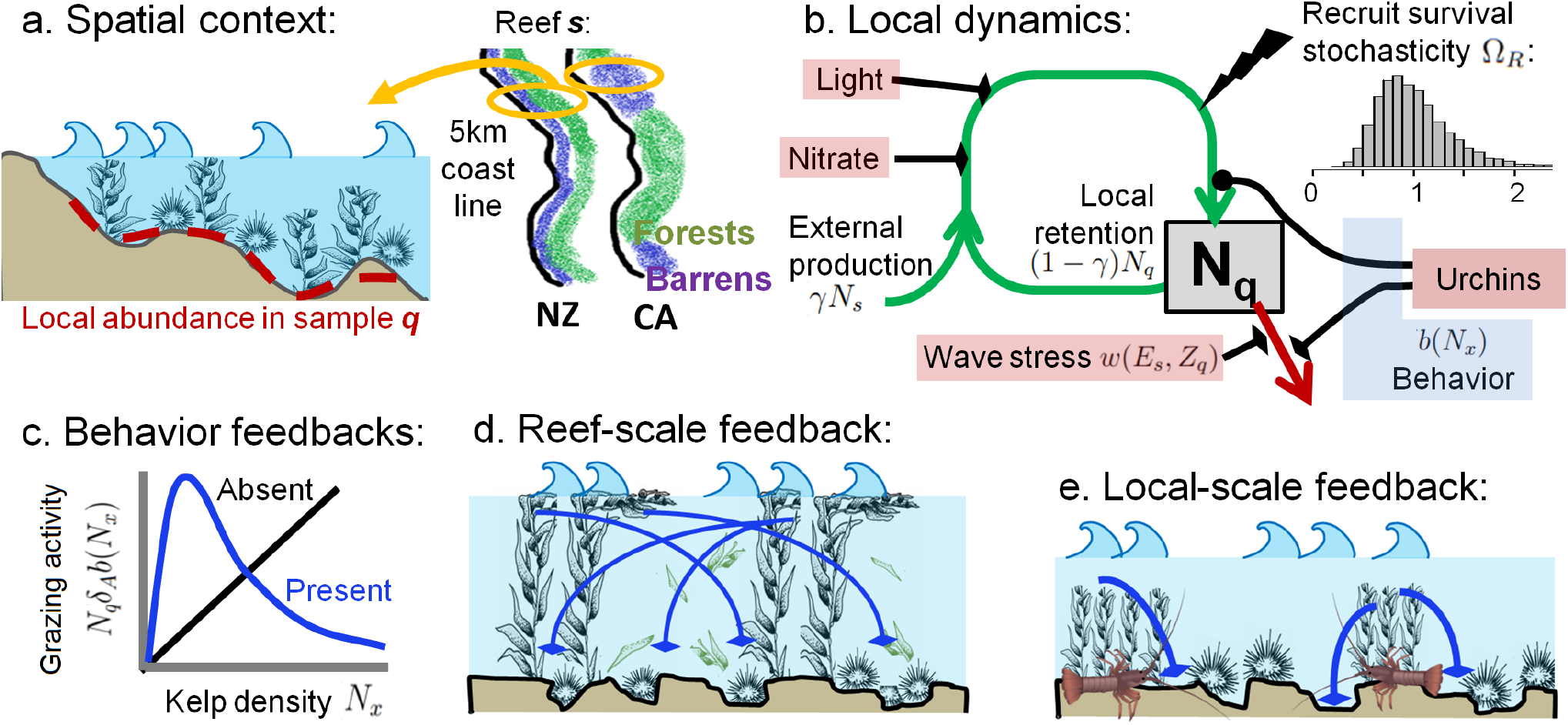
Model layout. (a) Local samples (red bars) on each reef span a depth gradient that influences wave intensity and light attenuation. Across reefs, forested and barren community types follow depth zonation in New Zealand but span entire reefs in California. (b) Dynamics of local kelp abundance *N*_*q*_ depend on environmental factors (red boxes), stochasticity, and urchin behavior (blue box) that affect adult survival (red line) or recruitment (green lines). Circular endpoints on lines denote negative effects and flat endpoints denote positive effects; stochasticity in recruit survival Ω_*R*_ (lightning bolt and inset distribution) can have positive (Ω_*R*_ > 1) or negative (Ω_*R*_ < 1) effects. (c) Functional form of grazing with behavioral feedbacks absent (black) or present (blue), where grazing rate declines with either local (*N*_*x*_ = *N*_*q*_) or reef-wide (*N*_*x*_ = *N*_*s*_) kelp density through urchin shifts from active to passive grazing. (d) Reef-scale feedbacks where passive drift kelp subsidies at high kelp densities *N*_*s*_ reduce urchin grazing across the reef (blue lines), as expected to be relevant in California. (e) Local-scale feedbacks where predators and physical abrasion in dense kelp stands deter grazing locally (blue lines), as expected to be relevant in New Zealand.

Here we evaluate whether or not behavior can affect large-scale community patterns in temperate rocky reefs by comparing observed patterns to predictions from models that incorporate environmental gradients, urchin density, and behavioral changes in urchin grazing. We represent behavioral changes in urchin grazing using a functional response with decreased grazing rate at high kelp densities, as might occur with a shift in grazing mode from active to passive with increased kelp density. The kelp density that affects grazing can be local as urchins avoid predators or abrasion or reef-wide as urchins passively graze on drift kelp. First, we test whether accounting for behavioral feedbacks better explains observed data, both compared to and in combination with environmental gradients. Second, we test whether the best-fitting behavioral feedbacks differ in spatial scale between regions and produce observed differences in patterning between New Zealand and California. Third, we test whether the best-fit models include alternative stable states, and if so, at what spatial scale.

## Methods

In this section, we first describe our systems data used to characterize kelp patterning and environmental drivers. Second, we describe our full model with environmental gradients, grazing, and behavior, from which we can exclude individual elements to explore processes alone or in combination. Third, we describe analyses of: (a) which factors best explain the data, (b) how incorporating behavior affects patterning in each region, and (c) whether best-fit models include alternative stable states.

### 2.1 Study systems

We focus our analysis on temperate rocky reefs in Northeast New Zealand (NZ) and the California Northern Channel Islands (CA) dominated by kelp (*Macrocystis pyrifera*, CA, and *Ecklonia radiata*, NZ) or urchins (*Strongylocentrotus purpuratus, Mesocentrotus franciscanus* in CA and *Evechinus chloroticus* in NZ). Like many temperate reefs, fast kelp growth and intense urchin grazing characterize these systems: abundant urchins can denude kelp forests in weeks, while under low urchin densities kelp can recolonize barrens within a few months (Ebeling *et al*. 1985). In contrast, urchin populations experience lower turnover and fluctuate more gradually in response to urchin predator abundance and multi-year changes in ocean climate (Shears *et al*. 2012; Okamoto 2014). This difference in time scales means that kelp abundance and urchin grazing activity can reach steady state under a given urchin density, whereas urchin density depends little on local kelp abundance due to demographic openness (Okamoto 2014) and can remain high in the absence of kelp (Filbee-Dexter & Scheibling 2014; Ling *et al*. 2015).

### 2.2 Data for kelp spatial patterns and environmental drivers

We use surveys of kelp density spanning 200-300km coastlines in each region, with 71 reefs sampled in 2001 in New Zealand (Shears & Babcock 2004) and 93 reefs sampled over 5-30 years in California (Kushner *et al*. 2013; Caselle *et al*. 2018). This geographic extent allows us to disentangle the effects of multiple processes by sampling a wide range of environments and increases robustness of our results to inter-annual variation in environment by exceeding the spatial scales of storms, upwelling variability, or recruitment pulses (8-50km; Cavanaugh *et al*. 2013; Karatayev & Baskett 2020).

Samples in both systems were collected at the end of the kelp growing season (July-August in CA, March-June in NZ) and span 100-500m^2^ of each reef *s* (see Appendix A for details on dataset and sampling methodologies). At the reef scale, we account for potential environmental drivers of kelp dynamics, including: total urchin predator density *P*_*s*_ (Shears *et al*. 2008; Caselle *et al*. 2018), wave stress *E*_*s*_ (Cavanaugh *et al*. 2011), and temperature-derived nitrate concentration *G*_*s*_ (Table S1). In the model below we omit nitrate limitation, which did not affect preliminary fits (Appendix A). Each reef was sampled using transects with 15-50 1-20m^2^ quadrat samples spanning a ∼ 15m water depth gradient (2,018 NZ samples, 16,539 CA samples). Within each quadrat *q*, we account for densities (ind m^−2^) of adult kelp plants and urchins (>25mm test diameter, *U*_*q*_), water depth *Z*_*q*_ (which attenuates waves), and near-bottom light availability *L*_*q*_. While *Ecklonia* can inhabit shallower depths than *Macrocystis*, in New Zealand we omitted 228 shallow (<3m) samples where wave-tolerant algae displaced both *Ecklonia* and urchins (Appendix A); we also omitted 777 samples along continuous transects that recorded *Ecklonia* presence but not density.

### 2.3 Model description

Our model follows dynamics of adult kelp abundance *N*_*q*_ in 1-5m^2^ locations *q* across a 0.1-0.5km^2^ reef *s* (Fig. 2a). We model kelp reproduction, spore survival, and adult survival as they depend on either local kelp density or kelp density averaged across the reef 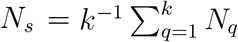 and reef-wide or local environmental factors. This model structure can produce kelp- and urchin-dominated regimes that form localized mosaics, gradients along depth-dependent environmental factors, or one regime spanning an entire reef. Below we describe the full model with all drivers, where zeroing out individual dynamics yields sub-models with different driver combinations.

Kelp reproduce continuously during the year with a baseline fecundity *m*. Newly produced spores can disperse over short distances (< 500*m* for *Macrocystis*, Anderson & North 1966; Reed *et al*. 1992). Therefore, we model the amount of spores *r* arriving in a location *q* as (1) the proportion *γ* of all spores dispersing throughout the reef plus (2) the proportion 1 − *γ* of locally-produced spores dispersing < 5*m*:

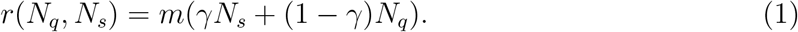

Survival of settled zygotes into adult sporophyte stages depends on both local and reef-scale factors. To incorporate the environmental gradient in light, we assume light availability increases with measured visibility *L*_*q*_ given proportionality constant *g*_*L*_ and declines with local density dominant adults *N*_*q*_ given proportionality constant *d* (Dayton 1985b). Due to their greater palatability compared with adults, juveniles also experience high rates of urchin grazing *δ*_*R*_ proportional to local urchin density *U*_*q*_, which can depend on behavior according to the function *b*(*N*_*x*_), as described below. We account for environmentally-driven stochasticity in survival, as might be due to thermal stress or interspecific competition, using a log-normally distributed random variable Ω_*R*_. Finally, given fast maturation (2-3 mo.) and high sensitivity of juveniles to competition and grazing, we assume juvenile abundance quickly reaches steady state on the time scale of adult kelp abundance, so that overall recruit survival is

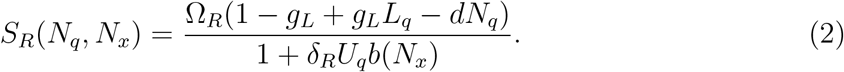

Adult mortality depends on local-scale urchin grazing and wave stress *E*_*s*_. To incorporate the environmental gradient in wave stress, we model wave stress dissipation by depth *Z*_*q*_ at more exposed sites following Bekkby *et al*. (2008). Dissipation depends on region-specific oceanographic features and weakens at exposed sites, represented by *f*_*w*_ and scaling factor *µ*. Per-capita adult mortality to wave stress is then

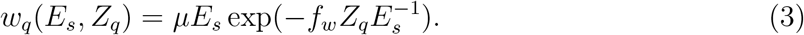

Grazing on adults occurs at a per-urchin rate *δ*_*A*_, as with recruit survival can depend on behavior according to *b*(*N*_*x*_) described below. Thus, per-capita adult mortality from grazing is

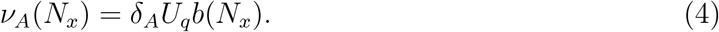

To incorporate a feedback between kelp density and urchin behavior in grazing, we assume urchin density is constant on the time scale of annual kelp dynamics but grazing rate can dynamically depend on adult kelp density (‘behavior feedbacks’ hereafter). Specifically, we incorporate the potential for a decline in grazing rate at high kelp densities, as might occur due to a switch from active to passive grazing mode, with a per-kelp grazing inhibition factor *ξ*_*A*_ in a generic ‘Type IV’ functional response (Koen-Alonso 2007; Bate & Hilker 2014). Dependence of this function on local-scale kelp density *N*_*x*_ = *N*_*q*_ represents localscale feedbacks involving physical abrasion (expected in NZ), and dependence on reef-scale kelp density *N*_*x*_ = *N*_*s*_ represents reef-scale feedbacks where drift kelp availability affects cryptic urchin behavior and associated direct grazing on kelp (expected in CA). Urchins may additionally avoid active grazing on reefs with high predator abundance *P*_*s*_. We integrate a per-predator grazing inhibition rate *ξ*_*P*_ over the entire year to arrive at the proportional decline in grazing due to predator avoidance, exp(−*ξ*_*P*_ *P*) (see Appendix B for separating predator effects on density *vs*. behavior). Altogether, behavior-mediated decline in urchin grazing is

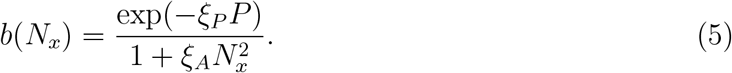

The overall dynamics of local kelp abundance are then

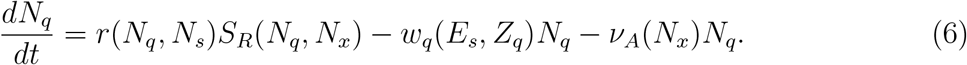

### 2.4 Role of urchin behavior

To compare the role of potential drivers (environmental, urchin density, behavioral), we compare the best fit of the full model (Eqn. 4) and simpler models that omit the effects of environmental factors (*µ* = *g*_*L*_ = 0), grazing (*δ*_*i*_ = 0), urchin predator avoidance (*ξ*_*P*_ = 0) and kelp-density feedback on urchin grazing rate (*ξ*_*A*_ = 0), each individually and in combinations. In Appendix C, we additionally evaluate whether herbivory dilution in a saturating Type II functional response, a behavior-independent feedback, can explain kelp patterning. We separate fits by region to estimate region-specific parameters. Given high kelp growth rates and our end-of-growing-season surveys (see “Study system”), we assume kelp densities reach steady state within one year. Therefore, we fit model equilibria to observed kelp abundance (i.e., a state-space model) and verify that best-fit models reach equilibrium within one year in Appendix D. Our approach assumes kelp densities at a reef are independent among consecutive years, an approximation we verify in Appendix E.

To evaluate model performance, we numerically solve models for the steady-state kelp abundance under the observed conditions (Table S1) in each sample and year, and then compare these predictions with observed kelp densities. We use a Runge-Kutta solver (R 3.4.3, deSolve package) with initial kelp abundance across the reef initially high or low (80th and 5th quantiles of observed densities in samples with kelp). Low initial abundance can occur when senescence pulses (in NZ) or storms (in CA; Cavanaugh *et al*. 2011) precede the growing season. To account for a range of possible recruitment conditions over the preceding year, we additionally solve the model under 10 realizations of Ω_*R*_ starting from each initial condition (i.e., 20 total realizations, Fig. 2c; Ω_*R*_ realizations identical across samples and models). To implicitly account for multiple secondary factors that allow kelp to occur in apparently adverse conditions (e.g., microsubstrate), we round predicted densities of 0 to 0.01.

For each quadrat we compute the Poisson probability of the observed kelp count under each realization, and average these probabilities across realizations to reflect that observed densities can arise from any combination of initial abundance and recruitment conditions (Gelman *et al*. 2013). We then calculate the logarithm of this marginal likelihood and, summing across all samples in the region, arrive at the total model log likelihood under a given parameter set. We find the best-fitting parameter sets using the DIRECT (global) followed by COBYLA (local) optimization algorithms in nloptr (Jones et al. 1993; Johnson 2019) and compare models based on Bayesian Information Criterion differences (ΔBIC), where ΔBIC>4 indicates improved model fit (Bolker 2008). We additionally calculate the squared correlation between predicted and observed (1) kelp densities 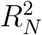 and (2) reef states (as defined below) 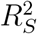 averaged across *S*_*R*_ realizations, choosing best-fitting initial conditions when models predicted two stable states.

As outlying kelp density observations favored maximal variation in Ω_*R*_, in the main analysis we set the Ω_*R*_ distribution mean to 1 and standard deviation to *σ*_*R*_=0.35 based on year-long recruitment experiments (Moreno & Sutherland 1982) (i.e., distribution parameters 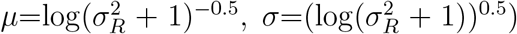. In Appendix E we evaluate model ranking robustness to ±30% changes in *σ*_*R*_, urchin-species-specific grazing, site effects, and temporal autocorrelation.

### 2.5 Drivers of community patterning and presence of alternative stable states

To resolve how grazing and wave stress gradients can jointly pattern communities, we use the best-fitting models from each region to project predicted kelp abundance over the observed range of reef-scale urchin densities. Within each reef we simulate 30 locations spanning the sampled depth range. Throughout, we set urchin distribution across depths and reef-scale wave stress, light availability, and predator density to the average values observed in data. We compare model projections with observed kelp patterns smoothed using 2-D splines.

To evaluate the role of behavior in explaining specific aspects of observed community patterns, we compare survey results with predictions of our best-fitting models with and without behavioral feedbacks. For this we first categorize the community state in each sample and model prediction (Fig. 1a, c). Given that urchins can maintain barrens even when rare (Ling *et al*. 2015), we classify samples and predictions with urchins and few kelp (CA: ≤ 0.05 individuals *m*^−2^, NZ: ≤ 1 individual *m*^−2^, 10th density quantiles in samples with kelp, Fig. 1a,b) as urchin-dominated, omit samples with no urchins and few kelp (NZ: 10% of samples, CA: 2%), and classify remaining samples as kelp-dominated. For within-reef patterning, we focus on a subset of reefs sampled by placing quadrats end-to-end in contiguous line transects (777 NZ samples, 5,644 CA samples) and quantify patch sizes as the number of adjacent samples with the same community regime.

## Results

### 3.1 Role of urchin behavior

Behavior-mediated grazing in California and an interactive effect of behavior-mediated grazing and environmental variation in New Zealand predominantly explain patterns in field data (Table 1). Best-fitting models in both regions include all three of environment, grazing, and behavior, and explain much of the variation in community states 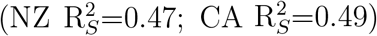. In models with behavior, local-scale grazing feedbacks (i.e., kelp-density-mediated grazing) in New Zealand and reef-scale grazing feedbacks in California best explain the data (Table 1). In California, models with grazing and behavioral feedbacks only outperform models with grazing and environmental factors only (ΔBIC=567). In both regions, models with behavior feedbacks outperform models without behavior because they can explain the co-occurrence of high and low kelp densities at intermediate urchin densities (Fig. 3). We find similar model ranking when controlling for higher red urchin grazing rates, temporal autocorrelation, site effects, and different levels of *σ*_*R*_ (Appendix E). We did not find support for Type II grazing saturation (Appendix C).

**Table 1:**
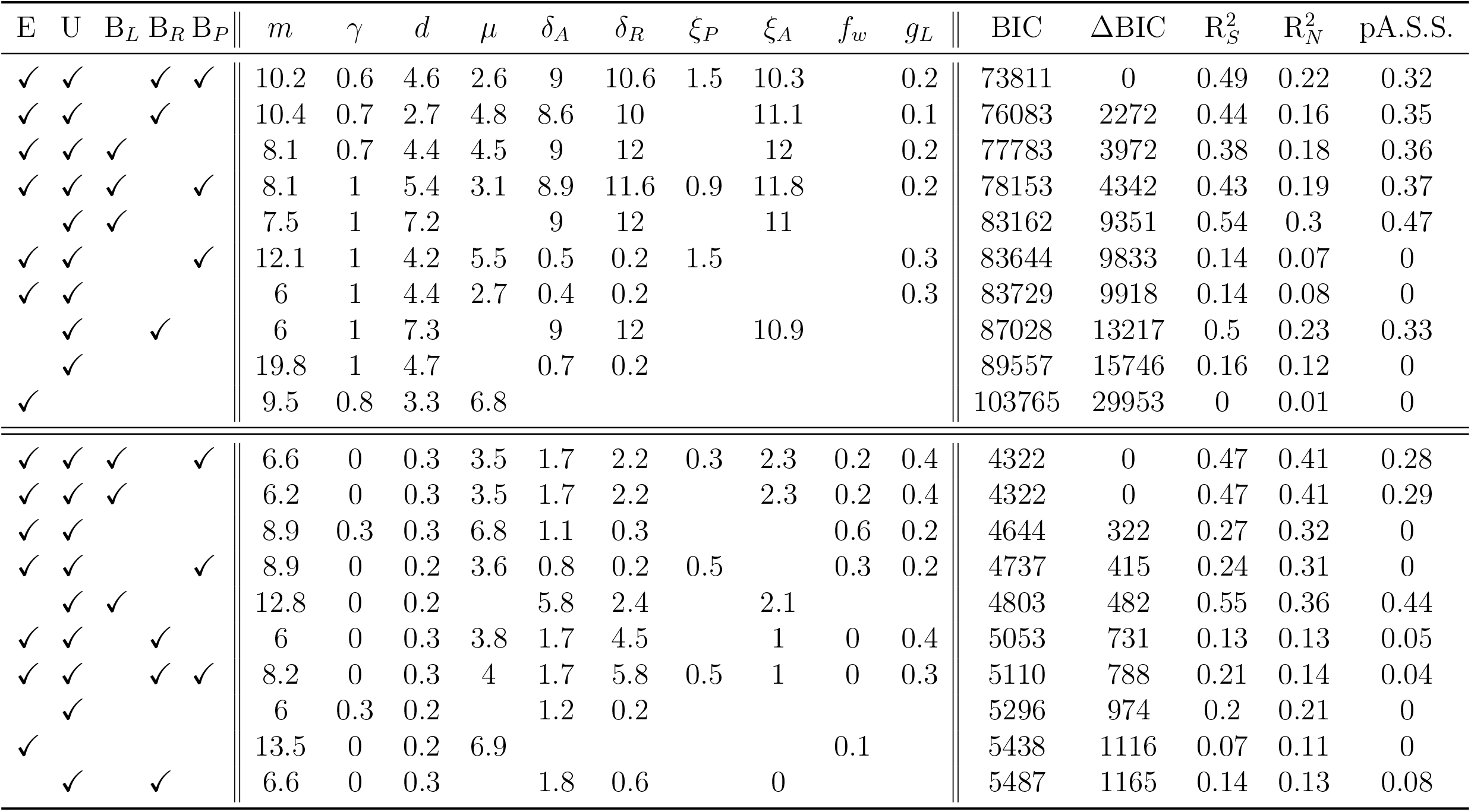
Results of model fitting and model comparison in California (top half) and New Zealand (bottom half). Check marks denote whether models include environmental factors (E), urchin grazing (U), local-scale behavior feedbacks (B_*L*_), reef-scale behavior feedbacks (B_*R*_), and predator avoidance (B_*P*_). pA.S.S. denotes the proportion of observations for which each model predicts alternative stable states. ΔBIC denotes BIC difference compared to the best-fitting model in each region. *R*^2^ columns denote squared correlation between observed and predicted reef states 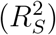 and kelp density 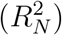. Covariate units are meters for *Z*_*q*_, and *L*_*q*_, *E*_*s*_, and *P*_*s*_ are proportions of their region-specific maximum values. See Appendix E for model-fitting details and sensitivity.

**Figure 3:**
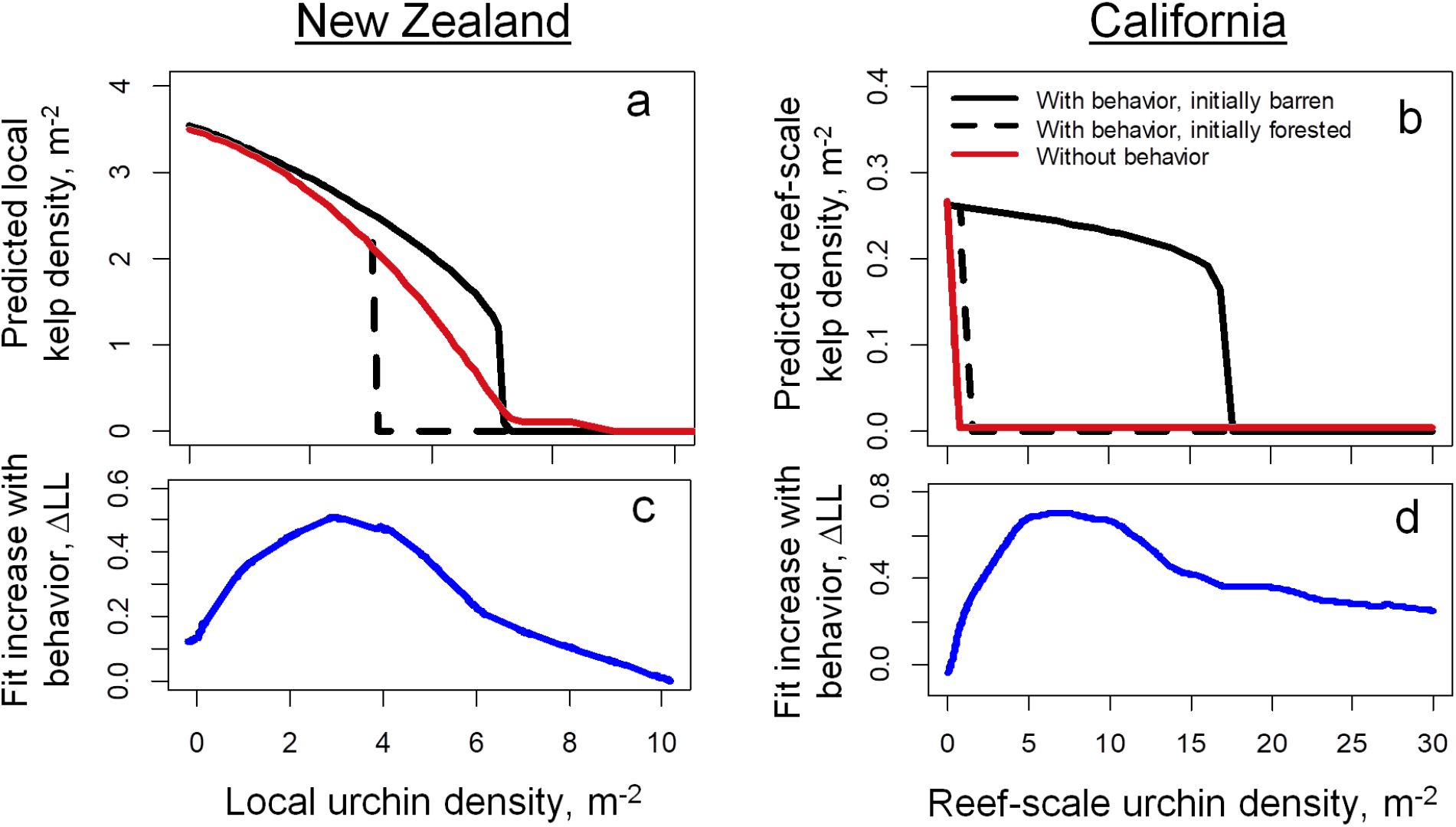
Behavioral feedbacks best explain observed patterns by predicting alternative stable kelp- and urchin-dominated states under moderate urchin densities. Here behavioral feedbacks occur through a decline in grazing rate at high kelp densities, as might occur thrugh a shift from active to passive grazing. (a, b) Kelp density predicted by best-fitting models without behavior (red lines) and models with behavior (black lines) for simulations with initially high (solid lines) and initially low kelp densities (dashed lines; without behavior, identical to the solid line). Note the comparison of local kelp densities in (a) and reefscale kelp densities in (b), reflecting the region-specific scale of feedbacks in our best-fitting models (Table 1). (c, d) Average difference in log likelihood between models with and without behavior. Note that lines in (a, b) denote deterministic equilibria under average environments; lower or higher levels of predators, visibility, and wave stress in specific samples broaden the range of urchin densities at which models with behavior predict alternative stable states and outperform models without behavior.

In both regions behavior improved model fit predominantly through kelp-density feed-backs as compared to predator-density-mediated behavior. Models with environment and kelp-density-mediated behavior outperformed models with environment and predator-density-mediated behavior (CA ΔBIC=7561; NZ ΔBIC=415). Nevertheless best-fit declines in grazing via predator avoidance appeared strong (up to 78% in CA, 26% in NZ) and strongly improved model fit in California (ΔBIC=2272). Lower support for predator avoidance in New Zealand possibly arose because few reefs had abundant predators or because predators congregate in kelp stands, an effect already captured by local kelp-density feedbacks.

### 3.2 Drivers of community patterning

The local scale of kelp-density behavioral feedbacks, combined with a decline in wave stress with depth, best explains the much smaller scale of community patterning in New Zealand than in California. In New Zealand, our best-fitting model predicts barrens as the only stable state in shallow areas (< 3*m*) due to a combination of grazing and high wave stress (Fig. 4b). At greater water depths that largely attenuate waves (> 8*m*), we predict forests are the only stable state because kelp quickly form dense stands that inhibit grazing. At intermediate wave stress between these zones, we predict that alternative stable states span 1-5m^2^ patches, where urchins concentrate grazing outside of dense kelp stands. This interface occurs at greater depths and urchin barrens cover a larger fraction of the system on reefs with greater overall urchin densities. In California, we predict that community regimes simultaneously span all reef depths due to the larger scale of grazing feedbacks (Fig. 4d, Table 1).

**Figure 4:**
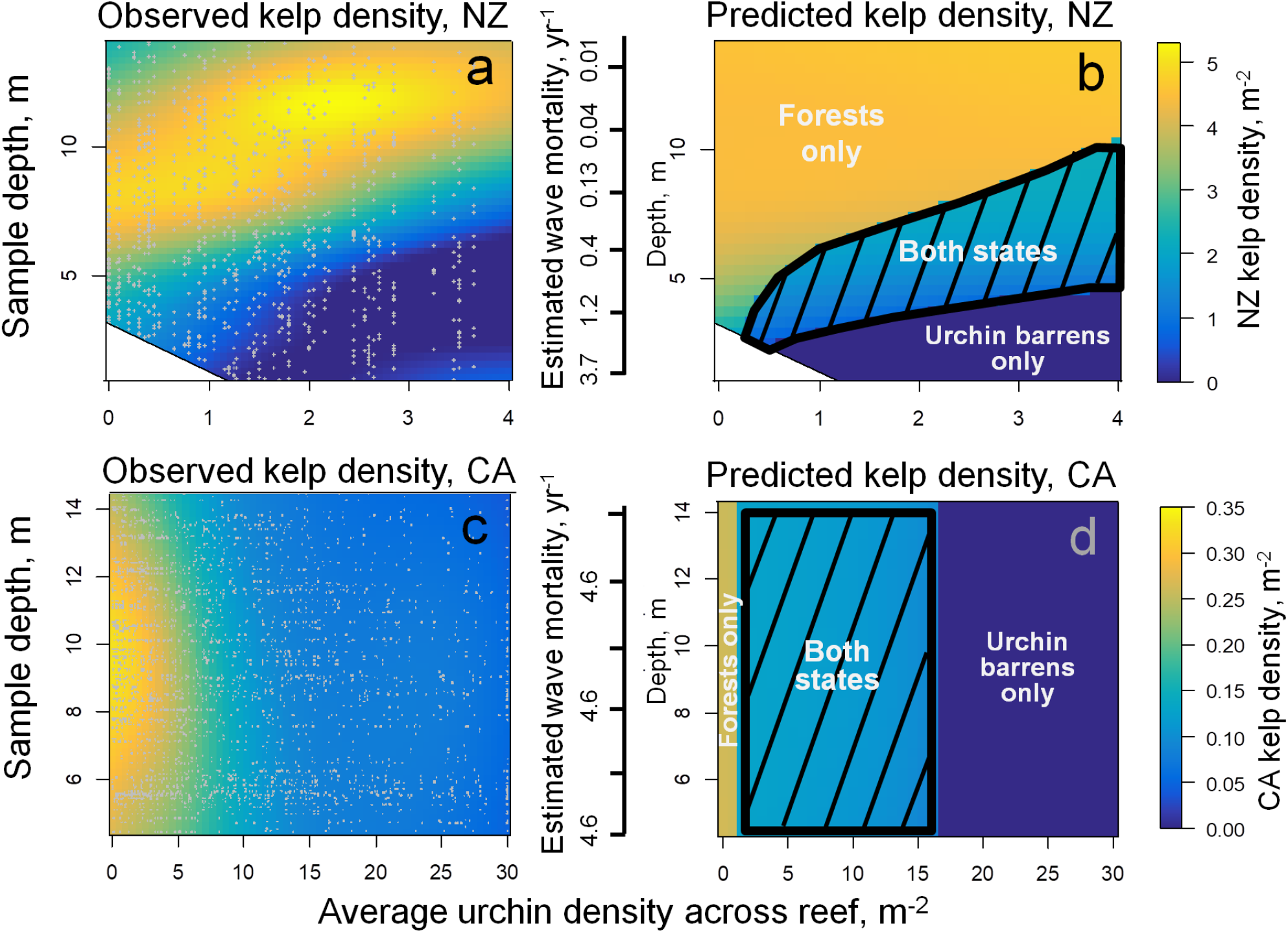
Regional differences in kelp distribution are best explained by a smaller scale of behavioral feedbacks and stronger wave stress gradients in New Zealand (a, b) compared to California (c, d). (a, c) Patterns in observed kelp density across depths on each reef (yaxes) and across reefs with increasing average urchin density. (b, d) Kelp densities predicted by best-fitting models in each region, with secondary y-axes denoting the best-fit, depthdependent estimates of mortality induced by wave stress. Gray dots in (a, c) denote the sample coverage across these conditions, with kelp density interpolated using 2-d splines. Hashed boxes in (b, d) denote conditions for which best-fitting models predict alternative stable states with kelp present or absent.

### 3.3 Presence of alternative stable states

Our best-fitting models predict that alternative stable states can occur in both regions (32% of CA samples, 28% of NZ samples, Figs. 3, 4b, d). In California, predicted alternative stable states occur at the scale of entire reefs (Fig. 5b, d). In New Zealand, alternative stable states occur over only a narrow range of depth-dependent wave stress intensities, such that our best-fitting model predicts that the final reef state depends little on initial kelp abundance (Fig. 5a). In this region, the small scale of kelp-and urchin-dominated patches observed in data arise only in models with local-scale behavior feedbacks (Fig. 5c). This shows that the spatial scale of behavioral feedbacks can determine the scale of patterning when feedbacks produce alternative stable states.

**Figure 5:**
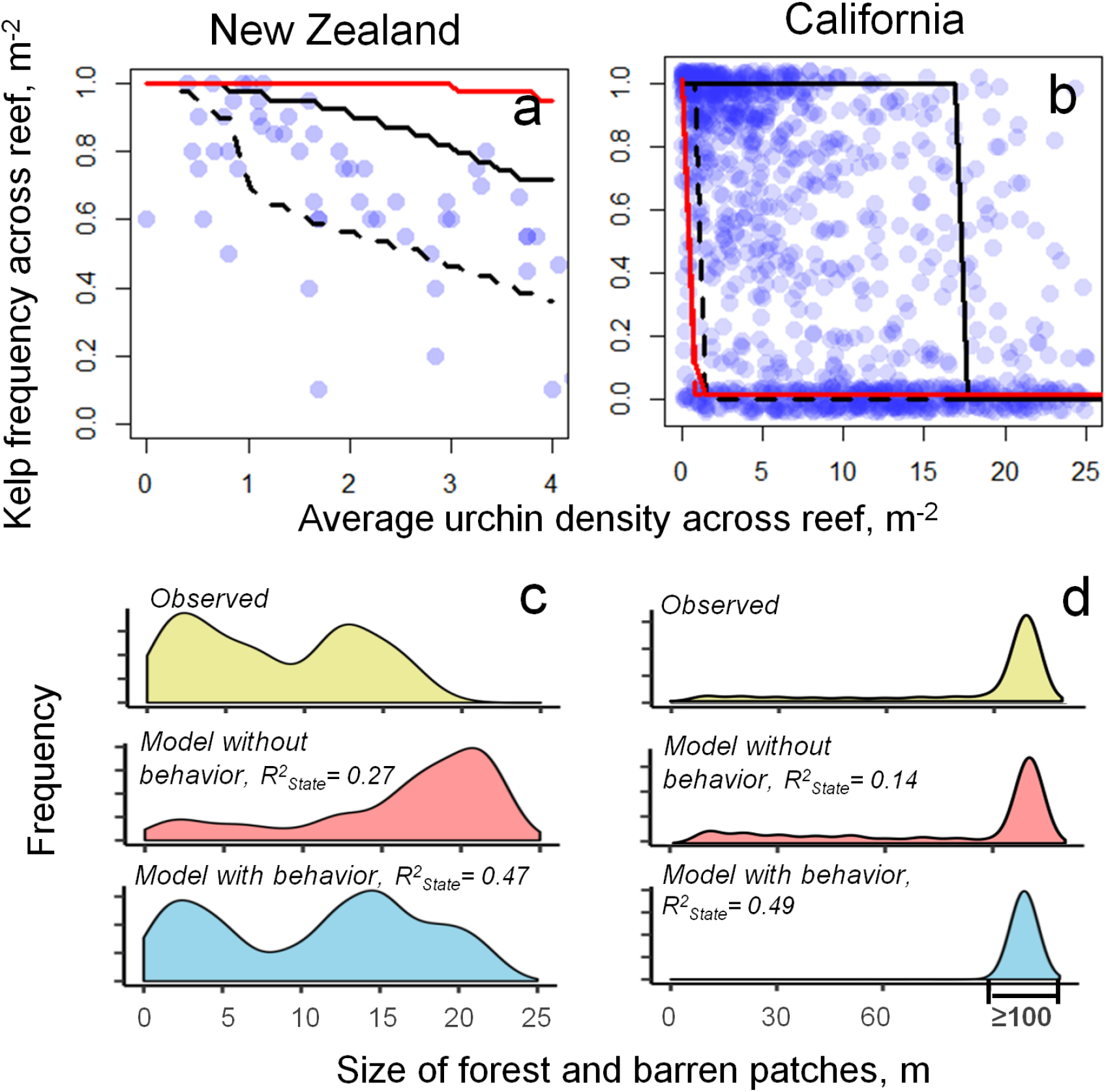
Best-fitting models with behavioral feedbacks predict alternative stable states that span a fraction of each reef in New Zealand (a) and entire reefs in California (b). (a, b) Frequency of kelp presence across reef predicted by best-fitting models without behavior (red lines) and models with behavior (black lines) for simulations with initially high (solid lines) and initially low kelp densities (dashed lines, identical to the solid line in the model without behavior). Blue dots show frequencies of kelp presence across all samples on each reef, with different dots representing different reefs and (in b) reefs in different years. (c,d) Sizes of barren and forested patches in data (yellow) and in best-fitting models without (red) and with (blue) behavior, with most patch sizes in California exceeding 100m due to limited spatial extent of sampling. 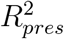 in (c, d) is the squared correlation between predicted and observed kelp presence (Table 1).

## Discussion

We show that behavior can determine the presence and scale of community patterning by mediating consumer-resource interactions in temperate rocky reefs. This potential occurs through feedbacks where kelp-density-dependent changes in urchin grazing rate amplify consumption when resources decline, as might occur through a shift from passive to active grazing modality. The scale of this feedback determines the spatial extent of the resulting resource- or consumer-dominated community regimes. Specifically, kelp forests and urchin barrens spanning entire reefs in California Channel Islands are most consistent with large-scale feedbacks, as might occur when drift kelp are transported over large distances and starvation-induced active grazing occurs when kelp densities decline across entire reefs (Figs. 4d, 5b; Table 1; Harrold & Reed 1985). In contrast, in Northeast New Zealand feedbacks that deter grazing locally, as might occur when kelp stands increase predation or physical abrasion (Ebeling *et al*. 1985; Konar 2000), can explain fine-scale patterning of community regimes organized into distinct depth zonation by a gradient in wave stress (Figs. 4b, 5a). Thus, feedbacks in consumer behavior interact with environmental heterogeneity to pattern communities at specific scales.

Our findings expand the results of prior behavior-induced patterning studies in two ways. First, our approach disentangles the role of grazer density from behavior by showing that models with environmental gradients and grazer densities do not predict observed patterns without also accounting for behavioral feedbacks (Table 1, Figs. 3, 5). Second, our best-supported models predict that behavioral feedbacks alone can create alternative stable states. Whenever localized pulse disturbances affect plant abundance (e.g., kelp loss in storms, Cavanaugh *et al*. 2011), this dynamic can produce persistent patterning in communities with spatially homogeneous environments and grazer densities. This extends results found in spatially heterogeneous systems, which show that behavior can mediate boundaries between community states (e.g., grazing halos, Matassa & Trussell 2011; Burkholder *et al*. 2013; Madin *et al*. 2019) but leave open the question of whether behavior feedbacks generate patchiness *de novo* in uniform environments.

While our results are built around temperate rocky reef systems, the Type IV functional response in urchin grazing behavior we include here can arise from the commonly-observed phenomena of starvation-induced consumption at low resource densities or group defense at high resource densities. Starvation-induced active urchin grazing might create persistent patterns observed in California because, after overgrazing kelp, urchins can survive for decades with little food due to low metabolic costs (Filbee-Dexter & Scheibling 2014; Ling *et al*. 2015). In other grazer taxa, however, starvation might not create persistent patterns if high metabolic costs cause starvation-induced mortality or grazers emigrate to higher-resource areas, and declining grazer densities allow eventual resource recovery. Group defense can arise through resource behavior when prey form schools or herds or, alternatively, through consumer behavior when herbivores avoid increased predation risk (Fortin *et al*. 2005) or environmental stress (Konar 2000) in dense vegetation. Persistent patterning of sparse and abundant resources, as in New Zealand reefs, can then arise when prey or plants in high-density patches exhibit group defense that shifts predation or herbivory to locations where resources are sparse (Schneider & Kefi 2016). As with our spatial model, Type IV functional responses from either factor can lead to alternative stable consumer- or resource-dominated states in a suite of spatially implicit models where consumer density depends on resource availability (Koen-Alonso 2007; Bate & Hilker 2014). Therefore, in addition to expanding empirical support for Type IV functional responses, our model suggests that Type IV functional responses might warrant exploration in other consumer-resource systems as a driver of spatial patterning.

### 4.1 Community patterning on temperate rocky reefs

While we fit models to New Zealand and California data, our findings can inform the drivers of community patterns in temperate rocky reefs globally. Mosaics separating urchin- and kelp-dominated depth zones found in New Zealand (Fig. 1, 4b, 5c) also occur in other systems dominated by sub-canopy kelp species, including northern Chile (Vásquez & Buschmann 1997), South Africa, (Ling *et al*. 2015), and Nova Scotia (Dayton 1985a). Compared to canopy-forming kelp, sub-canopy kelp might reduce long-distance drift subsidies through lower total biomass (here, 0.05kg m^−2^ in NZ *vs*. 2kg m^−2^ in CA; Shears & Babcock 2004; Cavanaugh *et al*. 2011) but increase short-distance urchin deterrence (e.g. ‘whiplash’, sheltering predators) as plants concentrate biomass near the bottom. Ubiquitous depth gradients in wave stress might also affect sub-canopy kelp more strongly than canopy-forming kelp because sub-canopy species can inhabit more exposed (<2m) depths while simultaneously being largely sheltered from wave stress in deeper (>10m) areas. However, kelp-urchin zonation may be reversed with barrens forming in deeper habitats when urchins are more sensitive to wave action than kelp (e.g., Chile, Nova Scotia; Dayton 1985a).

Large drift subsidies from canopy-forming kelp modeled here could underlie reef-scale grazing feedbacks and patchiness along the North American west coast. However, we expect behavior-driven patchiness to be less prevalent where grazing has weaker effects on kelp due to greater storm disturbance (e.g., central California, Cavanaugh *et al*. 2011), heat stress (southern and Baja California, Bell *et al*. 2018), urchin disease (Lafferty 2004), and predator densities (e.g., marine protected areas, Hamilton & Caselle 2015). Similarly, on *Macrocystis*-dominated reefs of central Chile, urchins rely primarily on passive grazing (Vásquez *et al*. 1984) but in other areas can form barrens. A second reef-scale feedback that can contribute to reef-scale forests and barrens arises when kelp facilitate recruitment of urchin predators (Karatayev & Baskett 2020), which freely forage across entire reefs (Topping *et al*. 2005). Our results highlight that behavior might strengthen this feedback in both regions as predators deter active grazing, complementing analogous findings in California (Ebeling *et al*. 1985; Caselle *et al*. 2018). Altogether, our results suggest that large-scale patchiness in systems characterized by canopy-forming kelp could arise from shifts in feeding modality among already-present urchins rather than urchin density changes via recruitment pulses and die-offs.

### 4.2 Model assumptions

To avoid model over-fitting, our approach leaves out additional potential dynamics that might drive alternative stable states and therefore patterning in temperate rocky reefs. First, we do not consider competition among primary producers which might displace competitively inferior juvenile stages of *Ecklonia* and *Macrocystis*, although <10% of samples indicate competitive exclusion by lacking both kelp and urchins (Fig. 1). Second, in our focus on whether or not behavior can explain observed patterns, we ignore many additional feedbacks hypothesized to drive (alone or in combination) alternative stable states in kelp forests (Ling *et al*. 2015). These additional feedbacks could increase the potential for patchiness. Further, we assume kelp abundance correlates with drift kelp that drives reef-scale grazing feedbacks; future biomass data may allow more mechanistic models of grazing. Finally, seasonality in wave-induced mortality (predominantly in winter) and recruitment (predominantly in spring) might weaken our assumption that kelp abundance reaches steady state within a year. Seasonal transients can obscure distinct equilibria (Mumby *et al*. 2013) and cause under-estimation of alternative stable states.

We also omit several secondary urchin behaviors that in both regions can increase the role of behavioral feedbacks in particular. Available data likely underestimates California urchin densities because cryptic urchins are harder to detect, causing best-fit models to underestimate the role of behavior in limiting grazing. Our model also omits urchin movement across the reef in response to kelp density, which can produce moving or stationary grazing fronts (Silliman *et al*. 2013) and in New Zealand might explain higher urchin densities in shallower areas. However, how kelp affects urchin movement in New Zealand is unclear because barrens do not expand following regular *Eklonia* senescence pulses. Given the strong role of behavior in community patterning found here, we suggest future experiments and more detailed spatial models explore the drivers of urchin movement.

### 4.3 Detection of alternative stable states

Behavioral feedbacks drive patchy spatial patterning in our best-supported models by giving rise to alternative stable states. Therefore, our results support the relevance of this phenomenon across both regions, especially in California where large-scale feedbacks can produce alternative stable states spanning entire reefs (Fig. 5b). In New Zealand, the localized scale of feedbacks limits alternative stable states to reef depths with intermediate wave stress on kelp (Fig. 4). These localized states average out to produce a gradual reef-wide response to changes in urchin density and little dependence of reef state on initial kelp abundance (Fig. 5a). This result supports existing theory (van Nes & Scheffer 2005) predicting greater relevance of alternative stable states at ecosystem scales either in spatially homogeneous environments or when biological feedbacks span large scales by involving mobile matter (e.g., drift kelp) or organisms.

The region-specific scales of alternative stable states found here can also help explain the debated presence of this phenomenon on temperate rocky reefs. Empirically demonstrating alternative stable states is challenging because of the limited spatiotemporal scales of experimental manipulations and measurements of biological feedbacks (Petraitis & Dudgeon 2004). Our results suggest one explanation for this debate: that the scale of alternative stable states is system-dependent. Future studies can estimate the potential scale of alternative stable states by quantifying the smallest observed areas of each state or the spatial scale of underlying ecological feedbacks.

Our analysis additionally expands on previous approaches to detecting alternative stable states. Studies most commonly test for this phenomenon based on whether distinct ecological states (e.g., Fig. 5b) or initial-condition dependency occur under the same levels of an environmental driver (Petraitis 2013; Mumby *et al*. 2013) by pooling observations or experiments across space or time. However, environmental heterogeneity may obscure distinct stable states (Mumby *et al*. 2013): for instance, our best-fit models predict that kelp densities in forested states double from 5 to 10m depths (Fig. 4b). Fitting dynamical models to time series can explicitly account for the expected effects of environmental variation (e.g., Ives *et al*. 2008), but requires long-term monitoring data. Instead, here we fit model steady states to data from large spatial surveys. Our analysis capitalizes on time scale differences between kelp abundance and comparatively slow changes in environment and urchin abundance, a feature utilized in previous kelp-urchin studies (Ling *et al*. 2015). We caution that this approach may produce biased interaction estimates by assuming population densities are independent among years. This approach additionally assumes that ecological interactions not modeled or measured explicitly vary little across locations and time; such variation can be accounted for using hierarchical modeling and model averaging techniques (Bolker 2008). Altogether, our work highlights how combining spatial surveys, mechanistic models, and statistics could predict the likelihood, given uncertainty and variable environments, that alternative stable states underlie observed ecological patterns.

## Acknowledgments

We would like to thank Alan Hastings, Jay Stachowicz, and three anonymous reviewers for feedback that improved the model and manuscript and Stephen Ellner and Stephan Munch for insights on model fitting. We also thank Tom Bell, Kevin Lafferty, and Katie Davis for access to data. Funding for this project was provided by the National Science Foundation Graduate Research Fellowship and the Davis-Auckland Graduate Student Exchange grant to VAK.

## Appendix

## Appendix A Dataset and covariates

We use survey data collected in New Zealand during the 1999-2000 winter growing season (Shears & Babcock 2004). For California, we use data collected in both the National Park Service Kelp Forest Monitoring survey (5644 samples, 1996-2017, ‘KFM’ hereafter; Kushner *et al*. 2013), which samples a wide range of years and collects many samples at each location, and the Partnership for the Interdisciplinary Studies of Coastal Oceans survey (9311 samples, 1999-2017, ‘PISCO’ hereafter; Casselle et al. 2015), which samples a greater number of reefs and a gradient of depths at each reef.

Adult kelp densities in New Zealand were sampled in 1m^2^ quadrats stratified by water depth. In California, kelp counts were sampled in either 20m^2^ areas (PISCO swath surveys) or in 5m^2^ quadrats (KFM); for KFM data we pooled data from adjacent quadrats to yeild kelp counts at the 20m^2^ scale. The difference in sampling scale across regions reflects the 20-fold lower densities of *Macrocystis pyrifera* in CA compared to *Ecklonia radiata* in NZ (Fig. 1). Analogously to New Zealand data, KFM surveys counted the abundance of > 1m tall adult *M. pyrifera* (plants with haptera at or above the primary dichotomy, Kushner *et al*. 2013). Based on KFM ‘Natural Habitat Size Frequency’ data for each reef and year, we found that the frequency of adults among > 1m tall *M. pyrifera* was best approximated by the frequency of > 1m tall plants with > 5 stipes 1m above the bottom (*R*^2^ = 0.57). Using this conversion, we determined the abundance of adult *M. pyrifera* in PISCO surveys, which quantified the number of stipes on each > 1 m tall *M. pyrifera* but did not classify plant life stage.

Urchin densities were quantified in stratified 1m^2^ quadrats (NZ), 20m^2^ areas (PISCO swath surveys) or in 1m^2^ quadrats (KFM). In all studies, density estimates represent urchins of sufficient size to consume kelp (≥ 25*mm* test diameter). As KFM surveys counted urchins of all sizes, we multiplied densities in KFM samples by the (species-specific) fraction of individuals ≥ 25mm in size-frequency data on ≥ 30 randomly selected individuals at each reef (‘Natural Habitat Size Frequencies’, Kushner *et al*. 2013). Additionally, as with kelp counts, for KFM data we averaged urchin densities among adjacent quadrats to estimate density (ind m^−^2) at the 20m^2^ scale. Finally, we pooled urchin density across purple (*Strongylocentrotus purpuratus*) and red (*Mesocentrotus franciscanus*) sea urchins in California samples.

In New Zealand, we omitted samples dominated by wave-tolerant brown algae which were not included in our model but can displace *Ecklonia* and sparse urchins from shallow areas. To systematically exclude these samples, we first identified samples dominated by brown algae (> 20 individuals, < 2 urchins, and < 5 *Ecklonia*, but results were not sensitive to these thresholds). To identify the depth extent of brown algal dominance, on each reef we calculated the 80th quantile of depths for samples dominated by brown algae. To account for the possibility that intense urchin grazing can exclude brown algae even from shallow areas, we fit a generalized additive model relating the depth extent of brown algal dominance to site-level urchin density using splines. This model predicted that brown algae are restricted to shallower depths at sites with more urchins. We then excluded the 228 samples where our model predicted brown algal dominance from our analysis (Fig. S1).

**Figure S1:**
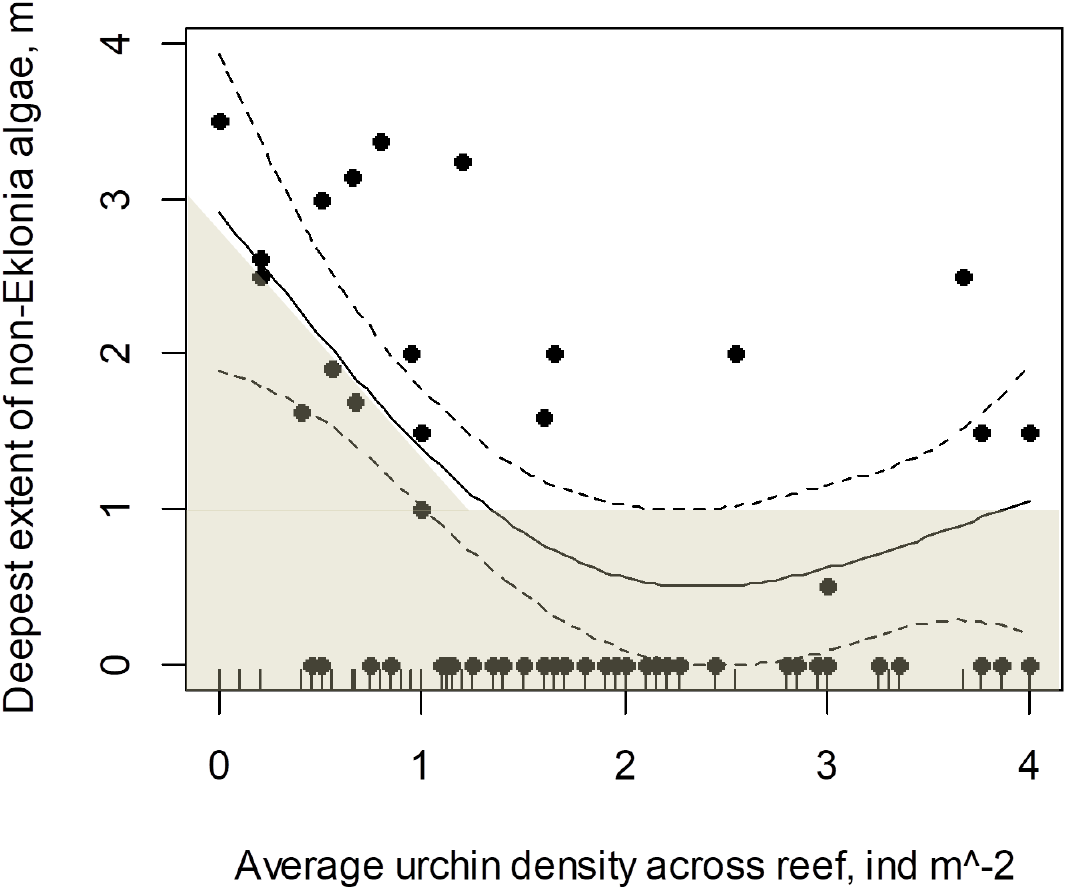
Depth extent of dominance by wave-tolerant brown algae (y-axis) on reefs in New Zealand (points) across sites with increasing urchin density (x-axis). Fitted curve represents a best-fit spline model, with dotted lines denoting model standard error. Shaded area denotes depths and site-level urchin densities at which samples were excluded from analysis.

For New Zealand, we determine predator density as the sum of snapper (*Pagrus auratus*) and lobster (*Jasus edwardsii*) densities estimated at the approximate time of kelp and urchin density surveys (Kelly et al. 2000; Willis et al. 2003; Shears *et al*. 2008). At KFM reefs in California, we determined predator density as the sum of sheephead (*Semicossyphus pulcher*) densities recored in Fish Visual Transects and sunflower seastar (*Pycnopodia helianthoides*) and spiny lobster (*Panulirus interruptus*) measured in 12 60*m*^2^ Band Transects (Kushner *et al*. 2013). At PISCO reefs in California, predator density is the sum of sheephead densities in fish visual surveys covering 120m^2^ (480m^3^ volume) and lobster and seastar densities in swath samples spanning 260m^2^ areas (mean swath area across reefs and years). Given greater sampling uncertainty in density (i.e., greater coefficient of variance) and their larger time scale of population dynamics compared with kelp and urchins, we smooth reef-scale predator densities at each site in California using a 3-year running average across sampling years.

We quantify the fraction of surface light reaching the bottom in each sample *q* at depth *Z*_*q*_ using reef-specific secchi depth measurements *D*_*s*_ as 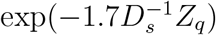. In New Zealand, wave stress *E*_*s*_ predominantly arises from waves generated by wind; we therefore use an index of potential wind fetch (Bekkby *et al*. 2008) as the proxy for wave stress. In California, kelp biomass can greatly depend on maximum wave height (Bell et al. 2015), but this environmental driver depends on both wind, coastal currents, and local reef topography. As a proxy for *E*_*s*_ we therefore use estimates of reef- and year-specific maximum wave height from (Lafferty et al. 2019), which calibrates maximum wave height estimates from regional oceanographic models using near-bottom sensors at each survey reef.

### 1.1 Effects of nitrate limitation

In California, large temperature variation driven by upwelling can cause nitrate depletion at high water temperatures (in New Zealand, lower peak temperatures rarely deplete nitrate). Nitrate limitation can reduce growth and total kelp biomass (not modeled here) as well as kelp fecundity (Bell *et al*. 2018). We include reef- and year-specific nitrate availability *G*_*s*_ estimated in Bell *et al*. 2018 from ocean temperature (projected locally using calibrated ocean circulation models) and observed relations of nitrate depletion with increasing temperatures. We modeled a decline in fecundity at low *G*_*s*_ using a function that saturates to 1 at high nitrate levels given saturation constant *v*, and change eqn. 1 to

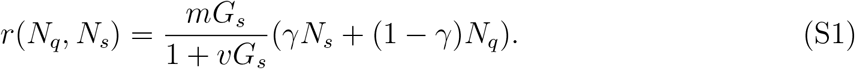

Adding nitrate to our models including environment, urchin grazing, and urchin behavior did not improve the fit of our best model based on BIC.

### 1.2 CA spatial patterning visualization

To illustrate spatial patterns of community states in Fig. 1d, we use satellite observations of kelp in 2016, during higher urchin densities were observed at all monitoring sites in the region (Fig. 1d, arrows). We classified all areas with kelp as forested and classified all areas without kelp in 2016 but where satellite imagery detected kelp in previous years as barren. To a limited extent this approach may over-estimate the spatial extent of barrens, for instance if storms cause kelp loss in some locations or as water currents temporarily push plants closer to the bottom, making detection more difficult. However, we do not expect such factors to affect patterning qualitatively (e.g., depth-independent patchiness).

---

**Table S1:** Model parameters fitted to data in each region and model covariates. Covariates are given by environmental data and parameters are estimated in model fitting. Covariates without units are set as proportions of region-specific maximum values to standardize parameter constraints.

| Parameter | Units | Description |
| --- | --- | --- |
| $m$ | $\text{yr}^{-1}$ | Adult kelp fecundity |
| $\sigma_R$ | | Standard deviation in recruit survival, fixed $\sigma_R = 0.35$ |
| $\gamma$ | | Fraction of kelp spores dispersing across reef |
| $g_L$ | | Contribution of measured light availability to recruitment |
| $d$ | $\text{N}^{-1}$ | Kelp competition for light inhibiting recruitment |
| $\mu$ | $\text{yr}^{-1}$ | Kelp mortality from wave stress |
| $f_w$ | $\text{m}^{-1}$ | Per-meter wave energy dissipation |
| $\delta_A$ | $\text{U}^{-1}\text{yr}^{-1}$ | Max grazing rate on adult kelp |
| $\delta_R$ | $\text{U}^{-1}\text{yr}^{-1}$ | Max grazing rate on kelp recruits |
| $\Gamma$ | $\text{N}^{-1}$ | Urchin grazing saturation constant |
| $\xi_A$ | $\text{N}^{-2}$ | Grazing inhibition by adult kelp |
| $\xi_P$ | | Grazing inhibition by predators |
| Covariate | Units | Description |
| $Z_q$ | m | Sample depth |
| $L_q$ | | Local light availability at bottom |
| $U_q$ | $\text{ind m}^{-2}$ | Local $> 25\text{mm}$ urchin density |
| $E_s$ | | Reef-scale wave stress |
| $G_s$ | | Reef-scale nitrate availability (CA only) |
| $P_s$ | | Reef-scale urchin predator density |

## Appendix B Behavioral declines in urchin grazing due to predators

Predators could affect grazing through both direct effects on urchin density (numerical effect) and indirect effects through changes in urchin grazing behavior (grazing activity effect); our model in the main text focuses on the latter. Examining the potential magnitude of numerical predator effects on urchins, we found that predator densities explain only a fraction of among-reef variation in urchin densities (averaged over all quadrats on each reef and sampling year, R^2^ =0.22, p<< 0.001, n=1267, Fig. S2a). Remarkably, the decline in urchin densities happened with only a small increase in predator densities (0 to 0.02 ind m^−2^), while further increases in predator density (0.02 to 0.15 ind m^−2^) had no effect on urchin density. Declines in urchin density over low predator densities only might also be explained by a strong urchin behavioral response where urchins are cryptic and harder to detect (i.e., quadrat samples under-estimating true densities) at sites where predators are present. To verify our density analysis, we examined whether predation reduced average size of urchins (test diameter of urchins < 70*mm*) found in quadrats along transects compared to urchins found in cages that largely exclude urchin predators (‘Artificial Recruitment Modules’, 5-15 permanent 5×12 cm mesh cages at each of 12 KFM monitoring sites). Overall, predator density poorly explained the difference in urchin size between quadrats and exclosures (R^2^ =0.12, p=0.0005, n=211, Fig. S2b), and urchins in exclosures did not have larger sizes at sites with high predator densities.

In New Zealand, large within-season changes in urchin density due to predation are unlikely because most sampled urchins were large and long-lived (Shears and Babcock 2004). Taken together, these results indicate that predators have only a limited direct, top-down effect on urchin densities. This reflects previous findings that predator presence strongly drives cryptic urchin behavior and reduced grazing activity and the fact that urchins comprise only a fraction (10-30%) of urchin predator diets. Note that, while predators could conceivably have species-specific effects on urchin predator avoidance *δ*_*U*_ (e.g., greater for slower-moving seastars), previous statistical analyses of patterns in the California Channel Islands did not detect species-specific *δ*_*U*_ (Caselle *et al*. 2018). We therefore approximate predator avoidance as dependent on total predator density only.

We additionally verify that data on kelp abundance (which reflects both total urchin abundance and per capita grazing activity) can distinguish the direct negative effects of predators on urchin density from indirect negative effects on urchin grazing activity. For this we construct a simple dynamical model of logistically growing urchin abundance *U* where both mortality and grazing activity *G* decline with predator density *P*. As with our base model, we assume that grazing activity has negligible effects on urchin density and that predator densities change little on the time scale of urchin population changes, yielding

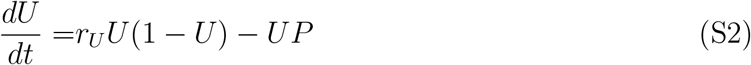

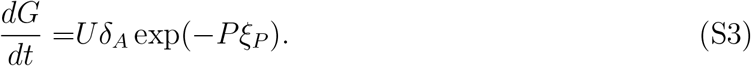

We then numerically solve this model for total grazing activity *U* at steady state (after 50-year transient) across levels of *P* in data (Fig. S4b) and the range of *ξ*_*P*_ estimates in our best-fit models. We find that predator avoidance (*ξ*_*P*_ > 0) distinctively reduces total urchin grazing compared to when grazing depends on urchin density only (Fig. S4c).

**Figure S2:**
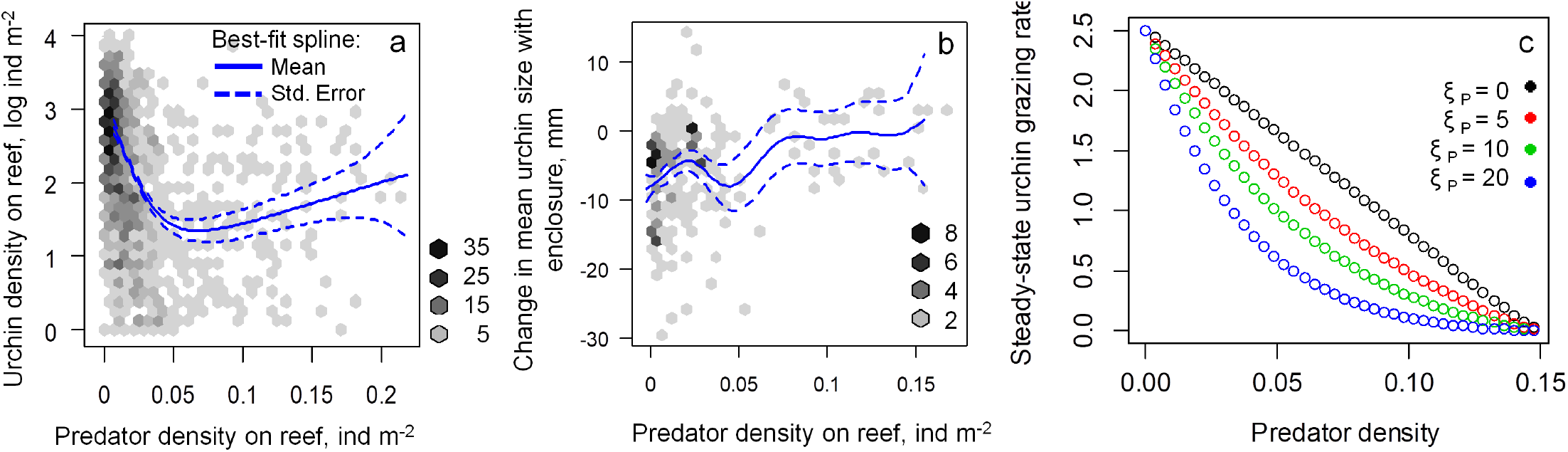
(a) Urchin densities correspond weakly with reef-scale predator densities and are characterized by lower urchin densities where predators occur, (b) mean sizes of 0-70mm urchins in predator exclosures rarely exceed mean urchin sizes outside predator exclosures, and (c) urchin grazing at steady-state without (*δ*_*U*_ = 0) and with (*δ*_*U*_ > 0) declines in urchin grazing activity with predator density. Blue lines in (a, b) denote best-fitting splines of predator density effects on each variable. Note that smaller urchin sizes in exclosures (negative values in b) at sites with few predators might arise from competition because urchin densities in exclosures were 1-2 fold greater compared to outside exclosures at all sites.

## Appendix C Role of herbivory dilution in patterning

Here we test whether herbivory dilution, a common behavior-independent kelp-density feedback (Noy-Meir 1975), can be an alternative explanation for observed kelp patterning. For this we evaluate models with a Type II functional response where grazing saturates with adult kelp density by a factor Γ, yielding an updated equation for recruit survival

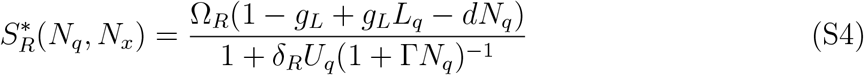

and adult mortality from grazing

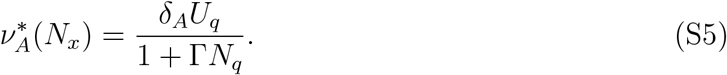

In fitting these models, we set the minimum constraint on *δ*_*A*_ to 0.01. We find that best-fitting models with behavior greatly outperformed models with grazing saturation in both regions (ΔBIC> 100, Table S2).

**Table S2:**
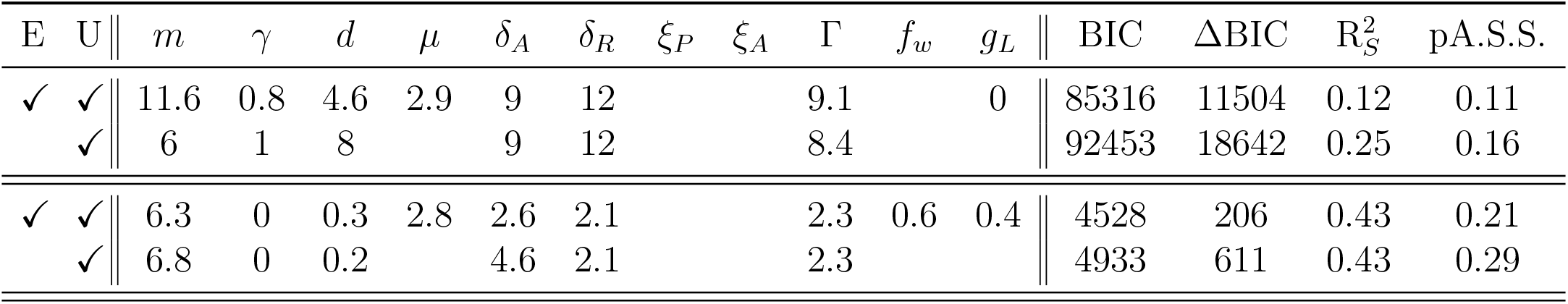
Results of model fitting and model comparison in California (bottom half) and New Zealand (top half) for models that include a Type II saturating grazing functional response and no behavior (i.e., assuming no subsidies, predator avoidance, abrasion, etc). ΔBIC denotes difference in fit compared to best-fitting models in each region in Table 1.

| E | U | $m$ | $\gamma$ | $d$ | $\mu$ | $\delta_A$ | $\delta_R$ | $\xi_P$ | $\xi_A$ | $\Gamma$ | $f_w$ | $g_L$ | BIC | $\Delta\text{BIC}$ | $R_S^2$ | pA.S.S. |
| --- | --- | --- | --- | --- | --- | --- | --- | --- | --- | --- | --- | --- | --- | --- | --- | --- |
| ✓ | ✓ | 11.6 | 0.8 | 4.6 | 2.9 | 9 | 12 |  |  | 9.1 |  | 0 | 85316 | 11504 | 0.12 | 0.11 |
|  | ✓ | 6 | 1 | 8 |  | 9 | 12 |  |  | 8.4 |  |  | 92453 | 18642 | 0.25 | 0.16 |
| ✓ | ✓ | 6.3 | 0 | 0.3 | 2.8 | 2.6 | 2.1 |  |  | 2.3 | 0.6 | 0.4 | 4528 | 206 | 0.43 | 0.21 |
|  | ✓ | 6.8 | 0 | 0.2 |  | 4.6 | 2.1 |  |  | 2.3 |  |  | 4933 | 611 | 0.43 | 0.29 |

## Appendix D Convergence deviations of fitted models

Here we determine the deviation of kelp abundance from model steady states after simulating population dynamics for a single year. We quantify these deviations for each model under the best-fit parameter set and across all initial conditions and realizations of juvenile survival stochasticity Ω_*R*_ (Table S3). In our best-supported models, kelp density approaches close to steady state within a single year in nearly all samples (kelp density difference of 99% in NZ, 93% in CA in samples with kelp; Fig. S3) and in most samples approaches close to steady state during the growing season (i.e., in ≤ 5 months, 98% in NZ, 83% in CA). All other fitted models analogously approach close to steady state within a year (Table S3). Greater deviations from equilibrium kelp abundance in California compared to New Zealand arise because the best-fit models in California assume strong coupling of kelp population dynamics across each reef by spore dispersal. This coupling among locations by recruitment leads to longer transient dynamics.

**Figure S3:**
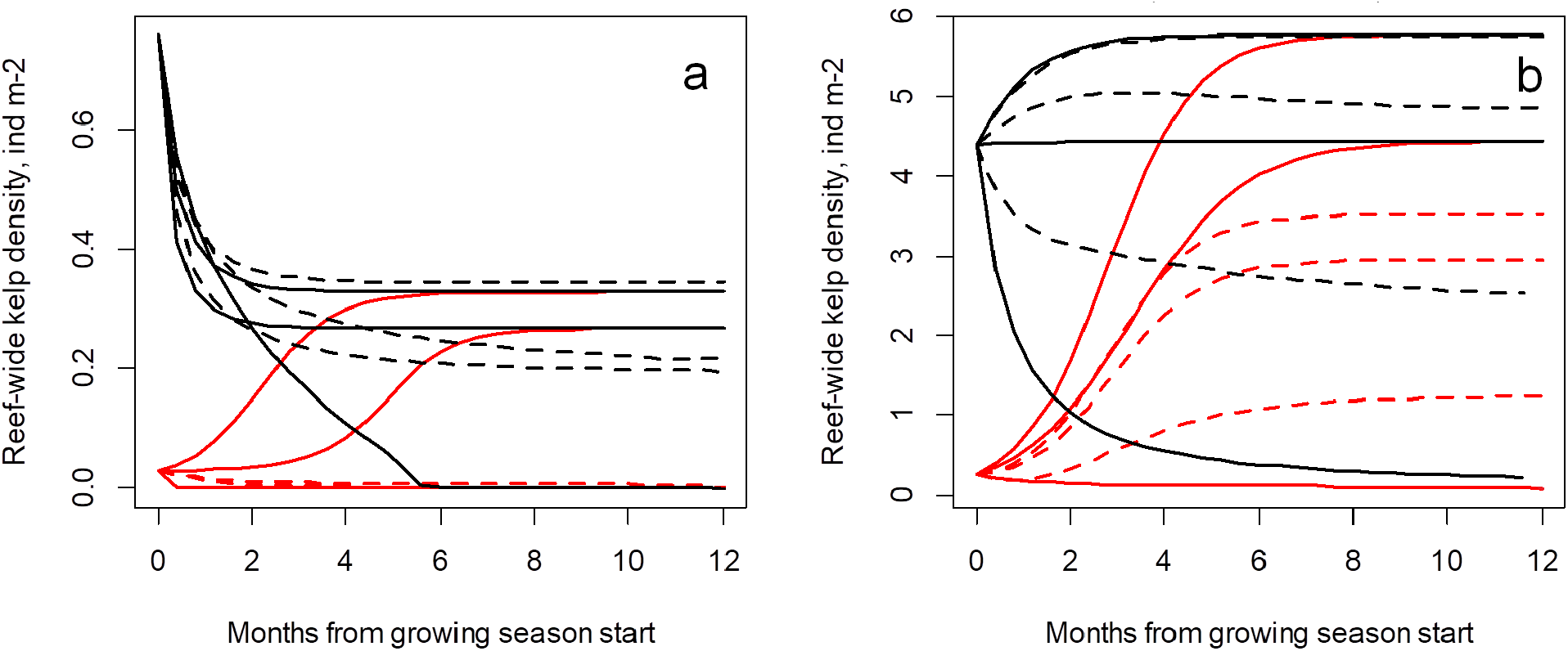
Within-year dynamics of kelp density (averaged over the entire reef) projected in best-supported models in California (a) and New Zealand (b). Trajectories in each system show dynamics from high (black lines) and low (red lines) initial kelp density for 3 (randomly selected) reefs with no alternative stable states predicted (solid lines), which converge on the same density, and 3 reefs with alternative stable states predicted (dashed lines).

**Table S3:** Mean and median deviation of kelp abundance from model steady states after simulating population dynamics for a single year, across all models fitted for New Zealand (top rows) and California (bottom rows). Relative (“Rel.” in table) deviations denote absolute deviations scaled by median kelp abundance in each region (4 in NZ, 0.2 in CA). Model nomenclature follows Table 1.

| E | U | B<sub>L</sub> | B<sub>R</sub> | B<sub>P</sub> | Mean Deviation | Mean Rel. Deviation | Median Rel. Deviation |
| --- | --- | --- | --- | --- | --- | --- | --- |
| ✓ | ✓ | ✓ |  | ✓ | 0.03 | 0.01 | 0 |
| ✓ | ✓ |  |  | ✓ | 0.1 | 0.03 | 0 |
| ✓ | ✓ |  |  |  | 0.05 | 0.01 | 0 |
| ✓ |  |  |  |  | 0.01 | 0 | 0 |
|  | ✓ | ✓ |  |  | 0.14 | 0.03 | 0 |
|  | ✓ |  |  |  | 0.11 | 0.03 | 0 |
|  | ✓ |  | ✓ |  | 0.13 | 0.03 | 0 |
| ✓ | ✓ |  | ✓ | ✓ | 0.03 | 0.13 | 0.01 |
| ✓ | ✓ | ✓ |  | ✓ | 0.03 | 0.14 | 0.01 |
|  | ✓ | ✓ |  |  | 0.02 | 0.08 | 0 |
|  | ✓ |  | ✓ |  | 0.02 | 0.11 | 0 |
| ✓ | ✓ |  |  | ✓ | 0.04 | 0.18 | 0.06 |
|  | ✓ |  |  |  | 0 | 0.02 | 0 |

## Appendix E Model-fitting details and sensitivity

### 5.1 Parameter constraints

We use several parameter constraints to ensure biologically plausible parameter fits (Table S4). We allow a level of urchin grazing on kelp recruits *δ*_*R*_ on par with levels of kelp population growth to reflect how low densities of urchins can prevent kelp recovery Ling *et al*. (2015). We set the maximum of grazing on adult kelp *δ*_*A*_ to be lower than that for *δ*_*R*_ because recruits are more vulnerable to urchin grazing than adults, which would require urchins to graze on stipes for kelp mortality to occur (Anderson et al. 1997). In New Zealand, we constrained maximum grazing inhibition by kelp to reflect that, due to their smaller size compared to *Macrocystis*, subcanopy kelp plants may be unable to deter urchin grazing fronts (Silliman *et al*. 2013), except possibly at high kelp densities. Finally, for wave stress dissipation *f*_*w*_ in California and grazing inhibition by predators *ξ*_*P*_ in both regions, we found that large maximum constraints encompassed local likelihood maxima that prevented optimizer convergence to better-fitting and biologically realistic parameters. To avoid these local maxima, we ran preliminary fits spanning a range of maximum parameter constraints on *f*_*w*_ and *ξ*_*P*_ and selected the weakest constraints (i.e., greatest possible maximum values) that permitted the best model fit. The resulting constrains on maximum grazing deterrence by site-level predator densities were lower in New Zealand compared to California, possibly reflecting that in NZ predators aggregate in dense kelp stands within each reef.

**Table S4:** Parameter constraints used in model-fitting. All parameters not listed were constrained only to positive values or proportions.

| Parameter | Region | Min | Initial | Max | Reference |
| --- | --- | --- | --- | --- | --- |
| $m$ | both | 6 | 8 | 20 | Baskett *et al.* 2007 |
| $\gamma$ | NZ | 0 | 0 | 0.3 | Itou *et al.* 2002 |
| $1/d$ | CA | 0.1 | 0.125 | 0.5 | 20<sup>th</sup> (min) and 80<sup>th</sup> (max) quantiles of regional kelp density in samples with kelp |
| $1/d$ | NZ | 1 | 2 | 6.7 | |
| $\mu$ | both | 0.001 | 1 | 7 | Cavanaugh *et al.* 2011 |
| $\delta_A$ | both | 0.25 | 2 | 9 | *see text* |
| $\delta_R$ | both | 0.25 | 10 | 12 | *see text* |
| $\log \xi_A$ | NZ | 0 | | 3 | *see text* |
| $\xi_P$ | CA | 0 | 0.15 | 1.5 | Shears *et al.* 2008; fitted, *see text* |
| $\xi_P$ | NZ | 0 | 0.15 | 0.6 | Shears *et al.* 2008; fitted, *see text* |
| $f_w$ | CA | 0 | 0 | 0.001 | fitted, *see text* |

### 5.2 Trends in model fit

Parsing the log likelihood of our best-fitting models across samples in each region, we find relatively consistent trends in model fit across space, time, and covariates, with two exceptions (Fig. S4). First, model fit worsens for sites farther west in the California Channel Islands; we attribute this trend to known environmental gradients that can change species interactions (Bonaviri et al. 2017) and the magnitude of environmental extremes. Second, in both regions model fit worsens at low urchin densities. As the more important driver of kelp density in both regions compared to environment (Table 1), low urchin densities often allowed kelp to reach carrying capacity in models. In data, however, peak kelp densities spanned a wide range of values across different sites (CA: 0.3-1.2 ind m^−2^, NZ: 4-16 ind m^−2^, ranges of 90th quantiles of kelp density across all samples with kelp at each site). We attribute this variation, and therefore reduced model fit at low urchin density, to site-specific environmental features not captured in our dataset (e.g., substrate composition or complexity). Note that reduced fit at low urchin density is also the likely explanation for changes in fit across light and wave stress in New Zealand due to the fact that, in this region, low urchin densities occurred at greater depths that corresponded to higher kelp densities, lower wave stress, and lower light availability.

**Figure S4:**
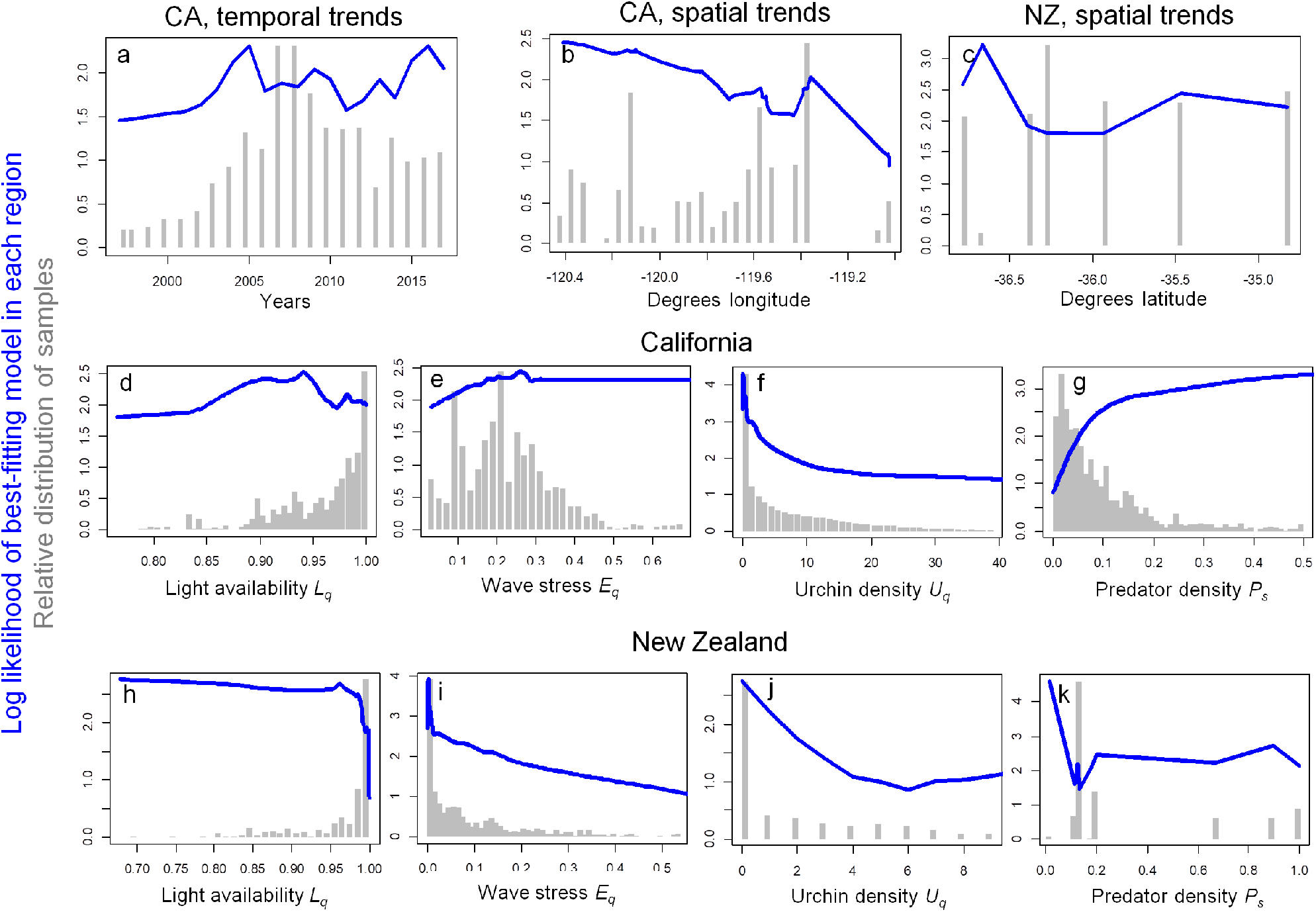
Trends in sample-specific log likelihood of best-fitting models in each region across years (a, CA only), space (b, c), and individual covariates (d-k). Y-axes denote the log likelihood of kelp density observed in each sample (i.e., as in our main analysis but before summing log likelihood over all observations in each region). In all panels, blue lines denote smoothed trends in sample-specific log likelihood and gray bars denote the relative distribution of samples across space, time, and covariates. Note that the spatial distribution of samples falls predominantly on an East-West orientation in California (spanning strong East-West environmental gradients in the Channel Islands; Kushner *et al*. 2013) and on a North-South orientation in New Zealand (Shears & Babcock 2004).

### 5.3 Spatiotemporal covariate autocorrelation and site effects

To verify that environmental autocorrelation does not qualitatively affect our results, we first calculate temporal and spatial autocorrelation for California data. Across site-level kelp densities and model covariates, these analyses show generally low (< 0.5) correlation even among adjacent sites and consecutive years (Fig. S5) with the exception of wave stress, which shows region-wide synchrony. To verify the robustness of our qualitative results on the importance of processes to temporal autocorrelation in California, we re-fit models to subsets of the full dataset that omit consecutive years from data at each site (i.e., using data from every other year, approximately 60% of CA observations). Model ranking in this analysis (Table S5) was analogous to that using the full dataset (Table 1). An exception to this result is that models with behavior via kelp-density feedbacks and predator avoidance perform worse than models with behavior via kelp-density feedbacks only. This might arise because most of the data omitted in this analysis come from long-term KFM monitoring sites, many of which are located in marine reserves.

To verify that site-specific features do not substantially skew our results, we re-fit models in both regions 20 sub-sets of our full dataset, each of which randomly selects 70% of all sites used in our base model fitting (we did not omit a larger portion of data to ensure models remained estimable). Across these replicate fits, models with kelp-density feedbacks in urchin behavior outperformed the best-fitting models without behavior (in 19 out of 20 replicate fits in CA and in all replicate fits in NZ). We also found relatively consistent parameter estimates across replicate fits (Table S6).

**Figure S5:**
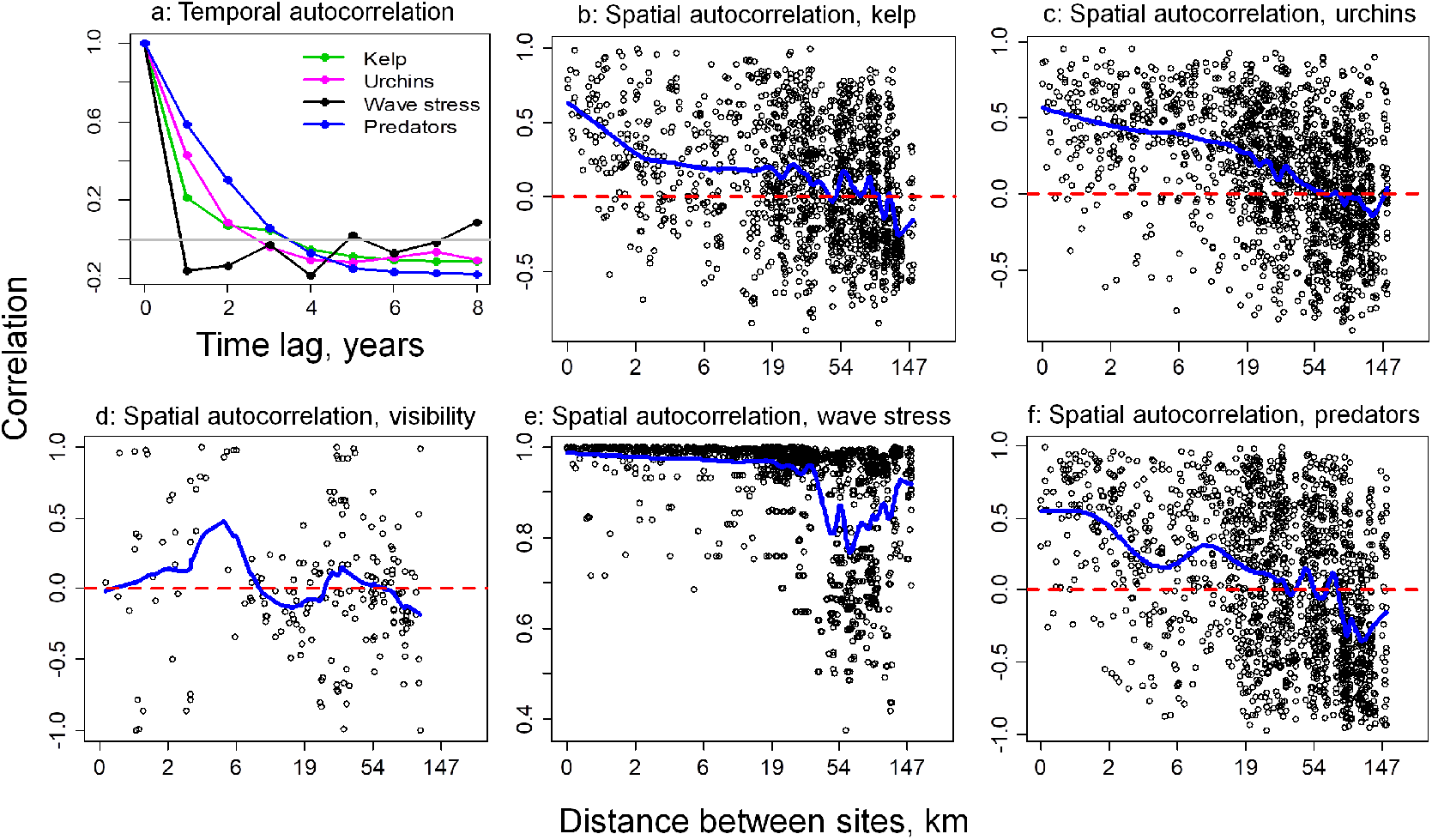
Correlation in site-level kelp densities and model covariates across years (a) and space (b-f) in California. Spatial autocorrelations are calculated using all sites with ≤ 5 years of data but omit comparisons between reefs which face opposite island faces (i.e., southern *vs*. northern) and therefore can experience distinctive oceanographic environments. In (b-f) blue lines denote smoothed running averages and axes are spaced on a log scale.

**Table S5:** Results of model fitting and model comparison in California omitting consecutive years from data at each site (i.e., using data from every other year, approximately 60% of CA observations).

| E | U | B<sub>L</sub> | B<sub>R</sub> | B<sub>P</sub> | $m$ | $\gamma$ | $d$ | $\mu$ | $\delta_A$ | $\delta_R$ | $\xi_P$ | $\xi_A$ | $f_w$ | $g_L$ | BIC | ΔBIC |
| --- | --- | --- | --- | --- | --- | --- | --- | --- | --- | --- | --- | --- | --- | --- | --- | --- |
| ✓ | ✓ |  | ✓ |  | 9.5 | 1 | 4.6 | 2.6 | 8.9 | 11.7 |  | 10.7 |  | 0.2 | 41807 | 0 |
| ✓ | ✓ |  | ✓ | ✓ | 9.5 | 1 | 4.7 | 2.6 | 8.8 | 11.6 |  | 10.7 |  | 0.1 | 41914 | 107 |
| ✓ | ✓ | ✓ |  | ✓ | 10.2 | 1 | 4.9 | 2.6 | 8.8 | 11.8 |  | 11.4 |  | 0.1 | 42090 | 283 |
| ✓ | ✓ | ✓ |  |  | 10.3 | 1 | 5.3 | 3.5 | 8.9 | 11.7 |  | 11.8 |  | 0.1 | 45150 | 3342 |
|  | ✓ | ✓ |  |  | 6.1 | 1 | 7.3 |  | 9 | 11.9 |  | 10.9 |  |  | 46449 | 4642 |
| ✓ | ✓ |  |  |  | 6.5 | 1 | 4.2 | 3.5 | 0.4 | 0.2 |  |  |  | 0.3 | 46893 | 5085 |
|  | ✓ |  | ✓ |  | 6 | 1 | 7.3 |  | 9 | 12 |  | 10.8 |  |  | 47276 | 5468 |
| ✓ | ✓ |  |  | ✓ | 18.8 | 1 | 5.7 | 7 | 0.9 | 0.2 | 1.4 |  |  | 0.4 | 47557 | 5749 |
|  | ✓ |  |  |  | 18.7 | 1 | 5.1 |  | 0.7 | 0.2 |  |  |  |  | 50662 | 8854 |
| ✓ |  |  |  |  | 11.3 | 1 | 4.8 | 5.9 |  |  |  |  |  |  | 60442 | 18635 |

**Table S6:**
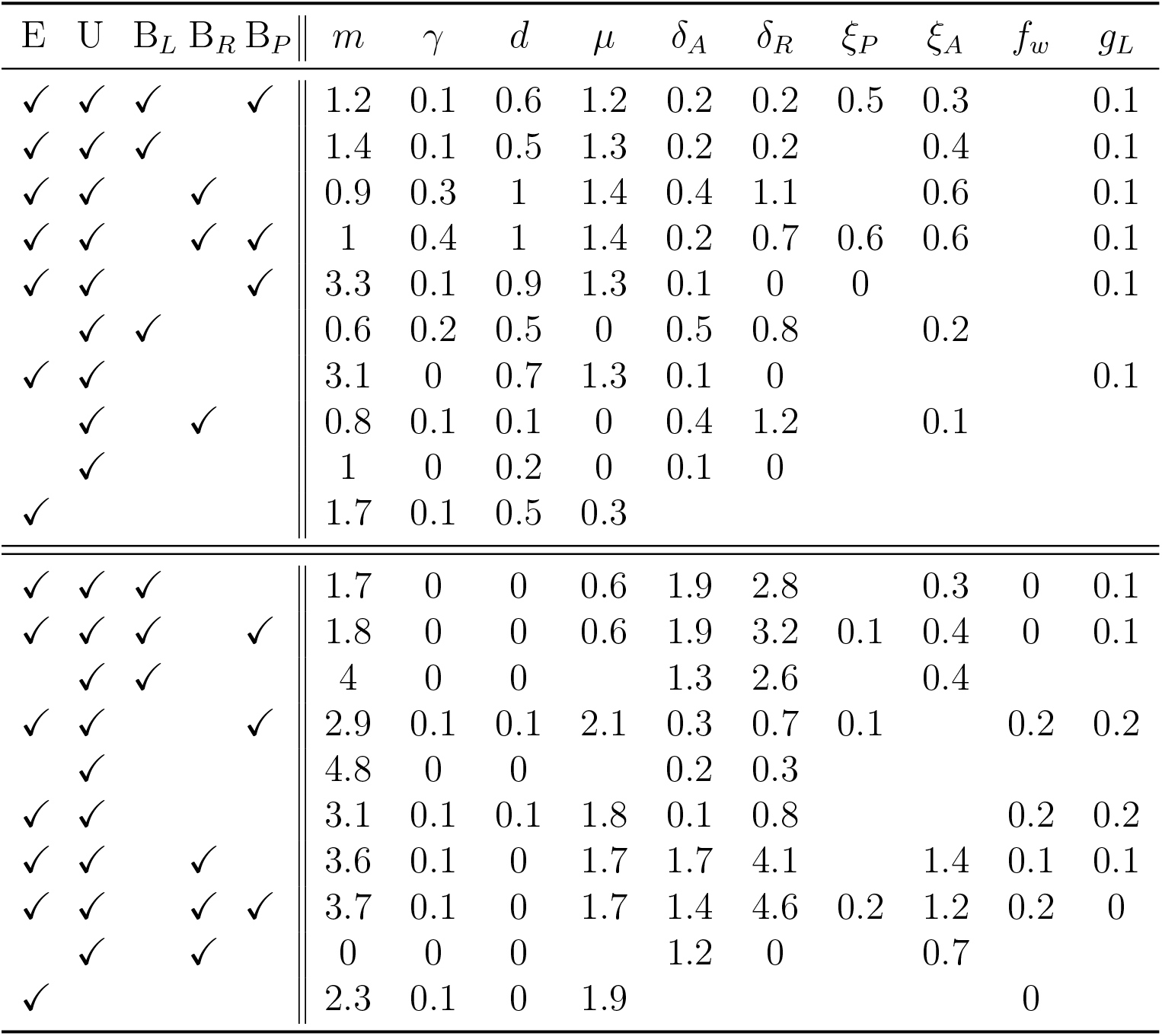
Standard deviations of fitted parameters across 20 replicate data sub-sets in California (bottom half) and New Zealand (top half). Each sub-set omits 30% of randomly selected sites in each region.

### 5.4 Species-specific urchin grazing in California

Our density-based approach to measuring urchin grazing does not account for the potential for larger urchins to graze at a higher rate (Stevenson et al. 2016). This could be a particular issue in California, where red urchins are substantially larger than purple urchins. Although we did not have data on the sizes of each urchin in our study, we explore the extent to which higher grazing rates by red urchins could affect our results. For this analysis, we first calculated the mean sizes of red and purple urchins measured at monitoring sites, converted urchin sizes to individual biomass and then to individual grazing rates following relations for *M. franciscanus* in Stevenson et al. (2016). We then calculated the ratios of per capita red *versus* purple urchin (size-based) grazing rates at each site and averaged these values across to arrive at a total estimate of ≈2.27-fold higher grazing rate for red compared to purple urchins. Using this ratio, we calculated a new index of grazing intensity 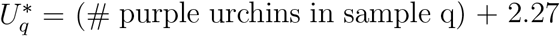 (# red urchins in sample q); for consistency of parameter estimates, we scaled 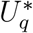 to have the same average value as *U*_*q*_ in our main analysis. Re-fitting our best models with and without behavior with using 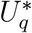 (Table S7), we find that accounting for higher grazing rates by red urchins does not affect model ranking but appears to reduce model fit compared to urchin density-based fits.

**Table S7:** Results of model-fitting with tests for sensitivity to species-specific urchin grazing in California (top rows), kelp recruitment stochasticity in California (middle rows), and kelp recruitment stochasticity in New Zealand (bottom rows).

| Test | E | U | B<sub>L</sub> | B<sub>R</sub> | B<sub>P</sub> | $m$ | $\gamma$ | $d$ | $\mu$ | $\delta_A$ | $\delta_R$ | $\xi_P$ | $\xi_A$ | $f_w$ | $g_L$ | BIC | pA.S.S. |
| --- | --- | --- | --- | --- | --- | --- | --- | --- | --- | --- | --- | --- | --- | --- | --- | --- | --- |
| U *spp.* | ✓ | ✓ |  | ✓ | ✓ | 10.7 | 1 | 4.6 | 5.1 | 8.4 | 11.7 | 1.1 | 11.7 |  | 0.2 | 79273 | 0.32 |
| U *spp.* | ✓ | ✓ |  | ✓ |  | 11.1 | 1 | 5.5 | 6.2 | 8.8 | 8 |  | 11.7 |  | 0.3 | 81869 | 0.3 |
| U *spp.* | ✓ | ✓ |  |  | ✓ | 13.2 | 0.8 | 4.4 | 6.5 | 0.4 | 0.2 | 1.5 |  |  | 0.3 | 83178 | 0 |
| $\sigma_R=0.45$ | ✓ | ✓ | | ✓ | ✓ | 9.7 | 0.3 | 2.8 | 4.7 | 8.8 | 9.8 | 1.3 | 11 | | 0.1 | 67995 | 0.27 |
| $\sigma_R=0.45$ | ✓ | ✓ | | | ✓ | 10.2 | 1 | 2.6 | 4.8 | 0.4 | 0.2 | 1.5 | | | 0.1 | 74133 | 0 |
| $\sigma_R=0.25$ | ✓ | ✓ | | ✓ | ✓ | 10.2 | 1 | 5.4 | 1.4 | 9 | 11.9 | 1.4 | 10.8 | | 0.1 | 80016 | 0.36 |
| $\sigma_R=0.25$ | ✓ | ✓ | | | ✓ | 14.2 | 1 | 5.7 | 4.3 | 0.6 | 0.2 | 1.3 | | | 0.3 | 88007 | 0 |
| $\sigma_R=0.45$ | ✓ | ✓ | ✓ | | ✓ | 13.3 | 0 | 0.3 | 1.9 | 2.9 | 6.7 | | 2.4 | 0.8 | 0.4 | 4665 | 0.37 |
| $\sigma_R=0.45$ | ✓ | ✓ | | | ✓ | 17.7 | 0 | 0.4 | 2.7 | 0.6 | 1 | 0.5 | | 0.8 | 0.3 | 5743 | 0 |
| $\sigma_R=0.25$ | ✓ | ✓ | ✓ | | ✓ | 14.3 | 0 | 0.3 | 2 | 5.3 | 5 | 0.5 | 2.9 | 0.7 | 0.2 | 4578 | 0.38 |
| $\sigma_R=0.25$ | ✓ | ✓ | | | ✓ | 16.2 | 0.3 | 0.2 | 7 | 0.7 | 1.7 | 0.5 | | 0.8 | 0.1 | 5006 | 0 |

### 5.5 Effects of recruitment stochasticity

To evaluate the sensitivity of model ranking to the magnitude of stochasticity in kelp recruitment *σ*_*R*_, we re-fit our best-fitting models with and without kelp-density feedbacks in urchin behavior with *σ*_*R*_ 30% higher and 30% lower than the base value used in our main analysis. Across both *σ*_*R*_ values and regions, models with kelp-density behavior feedbacks outperformed models without these feedbacks (Table S7). Parameter estimates were also generally similar to those in our main analysis (Table 1) with the exception of kelp competition *d*: with increasing *σ*_*R*_ best-fitting models estimate lower kelp competition to explain high kelp densities observed at some sites. This trend arises from our use of a lognormal distribution, where greater variation leads to a greater proportion of realizations with poor recruitment conditions.

## Notes

https://github.com/VadimKar/BehaviorPatternsCommunities

### Competing Interest Statement

The authors have declared no competing interest.

### Summary of Updates

Updated appendices, text, and figures for clarity.

